# Nanopore sequencing of *Giardia* reveals widespread intra-isolate structural variation

**DOI:** 10.1101/343541

**Authors:** Stephen M. J. Pollo, Sarah J. Reiling, Janneke Wit, Matthew L. Workentine, Rebecca A. Guy, G. William Batoff, Janet Yee, Brent R. Dixon, James D. Wasmuth

**Author notes:** Email Addresses (in same order as authors).

## Abstract

**Background:** Genomes of the parasite *Giardia duodenalis* are relatively small for eukaryotic genomes, yet there are only six publicly available. Difficulties in assembling the tetraploid *G. duodenalis* genome from short read sequencing data likely contribute to this lack of genomic information. We sequenced three isolates of *G. duodenalis* (AWB, BGS, and beaver) on the Oxford Nanopore Technologies MinION whose long reads have the potential to address genomic areas that are problematic for short reads.

**Results:** Using a hybrid approach that combines MinION long reads and Illumina short reads to take advantage of the continuity of the long reads and the accuracy of the short reads we generated reference quality genomes for each isolate. The genomes for two of the isolates were evaluated against the available reference genomes for comparison. The third genome for which there is no previous data was then assembled. The long reads were used to find structural variants in each isolate to examine heterozygosity. Consistent with previous findings based on SNPs, *Giardia* BGS was found to be considerably more heterozygous than the other isolates that are from Assemblage A. We also find an enrichment of variant-specific surface proteins in some of the structural variant regions.

**Conclusions:** Our results show that the MinION can be used to generate reference quality genomes in *Giardia* and further be used to identify structural variant regions that are an important source of genetic variation not previously examined in these parasites.

## Background

*Giardia duodenalis* (syn. *Giardia lamblia* or *Giardia intestinalis*) is a single-celled, eukaryotic, food and waterborne intestinal parasite that infects roughly 200 million people worldwide [1]. Infections can cause nausea, vomiting, diarrhea, and impaired growth and cognitive development [1]. The species *G. duodenalis* includes eight subtypes, named Assemblages A through H, at least two of which are known to infect humans (A and B) [1]. The cells have two diploid nuclei each containing five chromosome pairs [2]. The haploid genome size is ~12.8 MB [3]. Genome comparisons amongst assemblages of *G. duodenalis* found only 77% nucleotide and 78% amino acid identity in coding regions, suggesting the assemblages may represent different species [4]. Six isolates of *G. duodenalis* have reference genomes available [3].

Currently, whole genomes are sequenced using second generation technologies, third generation technologies, or strategies involving combinations of technologies (ex. combining PacBio and Illumina as in [5]). Second generation sequencing platforms produce high quality reads with low error rates (0.1% for Illumina HiSeq) but short lengths (mean length <250 bp for Illumina HiSeq), which pose challenges for assembly programs resulting in more fragmented assemblies [6]. In contrast, third generation sequencing platforms produce much longer reads (mean length <10 000 bp for PacBio and MinION) but have higher error rates (10-15% for PacBio and >10% for MinION depending on the chemistry) [6–8]. These longer reads have the potential to resolve many genomic areas that are problematic for second generation data, such as repetitive and/or duplicated regions [8]. Importantly, eukaryotic genomes have many such repetitive and duplicated regions (as much as two thirds of the human genome may be repetitive elements [9]), making eukaryotic genomes especially good candidates for sequencing with third generation technologies. Moreover, third generation data is well suited for examining structural variants within a genome. In diploid and polyploid organisms the different copies of each chromosome can contain large scale differences, including insertions, deletions, duplications, and translocations, in addition to variation at the single nucleotide level (SNPs). Collectively called structural variants, they are a major source of genetic variation, thought to play a larger role in phenotypic variation than SNPs, but are difficult to resolve using second generation data [10–12]. The tetraploidy of *Giardia* trophozoites further complicates short read genome assembly and structural variant detection methods because of the increased computational complexity of constructing four haplotypes for each locus. For a review on the challenges associated with polyploid eukaryotic genomes see [13]. Our expectation is that long read methods can detect and resolve the potentially three overlapping alternate alleles at any given locus.

The Oxford Nanopore Technologies (ONT) MinION is a third generation sequencing platform based on nanopore technology [8,14]. Briefly, the nucleic acids to be sequenced are driven through small pores in a membrane by an electrical current which causes fluctuations in the current in the pore [8]. Sensors measure these fluctuations, sending the data to a connected computer for processing and storage [8]. Assembling genomes *de novo* from MinION data involves basecalling of the squiggle files produced by the MinION during sequencing, assembly of the long reads into draft genomes, and polishing of the assemblies.

Here we have generated MinION and Illumina sequence data for *G. duodenalis* Assemblage A isolate WB (hereafter referred to as *Giardia* AWB), *G. duodenalis* Assemblage B isolate GS (hereafter referred to as *Giardia* BGS), and *G. duodenalis* isolated from a beaver (hereafter referred to as *Giardia* beaver). After generating reference quality assemblies with the long and short reads, the long reads produced here were then used to investigate heterozygosity in each isolate by detecting the structural variants in each genome.

## Data Description

We generated Oxford Nanopore Technologies MinION and Illumina MiSeq and iSeq whole genome sequence data for three isolates of *Giardia*. In addition to assembling genomes for the three isolates, we show the long read (MinION) data can be further used to detect structural variant regions within each genome. The sequences can be accessed from the sequence read archive (SRA) under accession number PRJNA561185.

## Analyses

### Reference quality assemblies

#### Performance of ONT long reads

The MinION sequencing runs used here produced several hundred thousand reads each with the exception of Run2, which was a second run conducted on a previously used flow cell (Table 1). In addition to producing fewer reads, re-using the flow cell also resulted in lower proportions of reads passing the quality threshold during basecalling with 64% and 81% of 1D reads passing in Run2 compared to 90 – 98% of 1D reads passing in Runs 1, 3, and 4 (Table 1). NanoOK [15] analysis of read error profiles showed that reads from Run2 have lower aligned base identity and higher substitutions per 100 bases compared to the other runs (Table 2).

**Table 1.** MinION sequencing run metadata, Albacore [28] basecalling results for both 1D and 1Dsq basecalling, and read statistics. “Pass” and “Fail” refer to reads that met or did not meet the quality threshold, respectively. Run 2 was conducted on a previously used flow cell after 64-72 h run time and so had few pores left.

| Name Used in this Document | AWB_0150 | AWB_0157 | AWB_2331 | AWB_2338 | Beaver_2302 | Beaver_2309 | BGS_2237 | BGS_2244 |
| --- | --- | --- | --- | --- | --- | --- | --- | --- |
| Run Name | SRRun1 | SRRun1 | SRRun2 | SRRun2 | SRRun3 | SRRun3 | SRRun4 | SRRun4 |
| Run ID | 20170720_0150_GiardiaWB_20170719 | 20170720_0157_GiardiaWB_20170719 | 20170721_2331_GiardiaWB_20170721 | 20170721_2338_GiardiaWB_20170721 | 20170726_2302_GiardiaBeaver_20170726 | 20170726_2309_GiardiaBeaver_20170726 | 20170731_2237_GiardiaGS_20170731 | 20170731_2244_GiardiaGS_20170731 |
| Isolate | *Giardia* AWB | *Giardia* AWB | *Giardia* AWB | *Giardia* AWB | *Giardia* beaver | *Giardia* beaver | *Giardia* BGS | *Giardia* BGS |
| Reference Genome Size (bp) | 12827416 | 12827416 | 12827416 | 12827416 | N/A | N/A | 11001532 | 11001532 |
| Total Number of 1D Reads | 1225 | 329039 | 237 | 19531 | 1668 | 382740 | 1508 | 885046 |
| Number of 1D Reads Pass | 1207 | 304219 | 152 | 15842 | 1603 | 354581 | 1449 | 804942 |
| Number of 1D Reads Fail | 18 | 24820 | 85 | 3689 | 65 | 28159 | 59 | 80104 |
| Total Number of 1Dsqs Reads | 172 | 60156 | 16 | 1904 | 146 | 53553 | 212 | 143371 |
| Number of 1Dsqs Reads Pass | 68 | 25755 | 0 | 192 | 69 | 29349 | 124 | 62452 |
| Number of 1Dsqs Reads Fail | 104 | 34401 | 16 | 1712 | 77 | 24204 | 88 | 80919 |
| Average Length of 1D Reads | 5066.15 | 7195.29 | 3450.08 | 6484.00 | 5113.00 | 8270.88 | 6534.03 | 9417.60 |
| Longest 1D Read | 42781 | 470735 | 32138 | 330795 | 37229 | 1132445 | 56642 | 485807 |
| Average Length of 1Dsqs Reads | 5335.22 | 7685.61 | 2853.62 | 7344.74 | 5273.86 | 8472.84 | 5529.57 | 9829.82 |
| Longest 1Dsqs Read | 18489 | 43102 | 6523 | 32705 | 22740 | 59564 | 25876 | 66185 |

**Table 2.** Read error profiles for *Giardia* AWB and *Giardia* BGS MinION sequencing runs.

| Error Type | AWB_01<br>50 Reads | AWB_01<br>57 Reads | AWB_23<br>31 Reads | AWB_23<br>38 Reads | BGS_22<br>37 Reads | BGS_22<br>44 Reads |
| --- | --- | --- | --- | --- | --- | --- |
| **Proportion of Reads Counted (%)** | 87.55 | 83.56 | 28.04 | 52.61 | 12.62 | 77.47 |
| **Overall Base Identity (%)** | 76.907 | 74.577 | 54.293 | 65.904 | 58.255 | 56.636 |
| **Aligned Base Identity (%)** | 90.526 | 89.352 | 83.076 | 83.915 | 91.429 | 89.954 |
| **Identical Bases per 100** | 80.430 | 78.338 | 71.024 | 71.597 | 80.855 | 78.834 |
| **Inserted Bases per 100** | 5.291 | 3.881 | 7.811 | 5.087 | 3.473 | 4.478 |
| **Deleted Bases per 100** | 5.860 | 8.450 | 6.758 | 9.592 | 8.105 | 7.886 |
| **Substitutions per 100** | 8.415 | 9.334 | 14.406 | 13.725 | 7.569 | 8.801 |
| **Mean Insertion** | 1.638 | 1.462 | 1.755 | 1.480 | 1.482 | 1.530 |
| **Mean Deletion** | 1.621 | 1.787 | 1.591 | 1.788 | 1.848 | 1.898 |

NanoOK analysis of 1D read error profiles for all runs indicated a 9 – 17% error rate in the regions of reads that aligned to the reference genome (Table 2, aligned base identity) and a 24 – 46% error rate across the entirety of reads that aligned to the reference genome (Table 2, overall base identity). The analysis also showed more deleted bases than inserted bases in the reads (Table 2). Average and maximum read lengths for all runs are presented in Table 1. Notably, the maximum 1D read length generated in the sequencing runs analyzed here was 1,132,445 bases, though this read did not align to any *Giardia* reference genome nor did it have significant BLAST hits longer than ~45 bp in the nr database (data not shown). It is presumably a strand that got stuck but continued to generate (incorrect) sequence data.

Of the 39 long read *de novo* assemblies performed (13 input combinations × 3 assembly programs; see Materials and Methods long read assembly evaluation), five did not have sufficient numbers of reads to generate any contigs (AWB_2338_1D_smartdenovo, AWB_2338_1Dsq for all three assemblers, and AWB_2331_2338_1D_smartdenovo). The remaining assemblies were all polished with Nanopolish eight times and the evaluation metrics were calculated for the nine resulting draft assemblies from each *Giardia* AWB and BGS input/assembler combination for a total of 315 assemblies (Supplementary Table 1). The top performing AWB and BGS assemblies for each metric are listed in Supplementary Table S2. No assembly ranked first in more than two of the metrics. To further examine the effects of 1D vs 1Dsq input reads, pooling reads for the same isolate from multiple runs, assembly program, and number of genome polishing iterations, for each metric the values for all the assemblies were plotted (Supplementary Figs. S1 – S10). The average value and standard deviation for each group were also calculated (Supplementary Tables S3 – S10). Figure 1 shows the effects of 1D vs 1Dsq input reads, assembly program, and number of genome polishing iterations on BGS assemblies for four of the metrics – the two that don’t require a reference genome (number of contigs and genome size), gene finding (BUSCO score), and accuracy measured as average percent identity. The averages and standard deviations that correspond to Figure 1 can be found in Supplementary Tables S4, S8, and S10. The other metrics and the values for AWB assemblies show similar trends (Supplementary Figs. S1 – S10).

**Figure 1.**
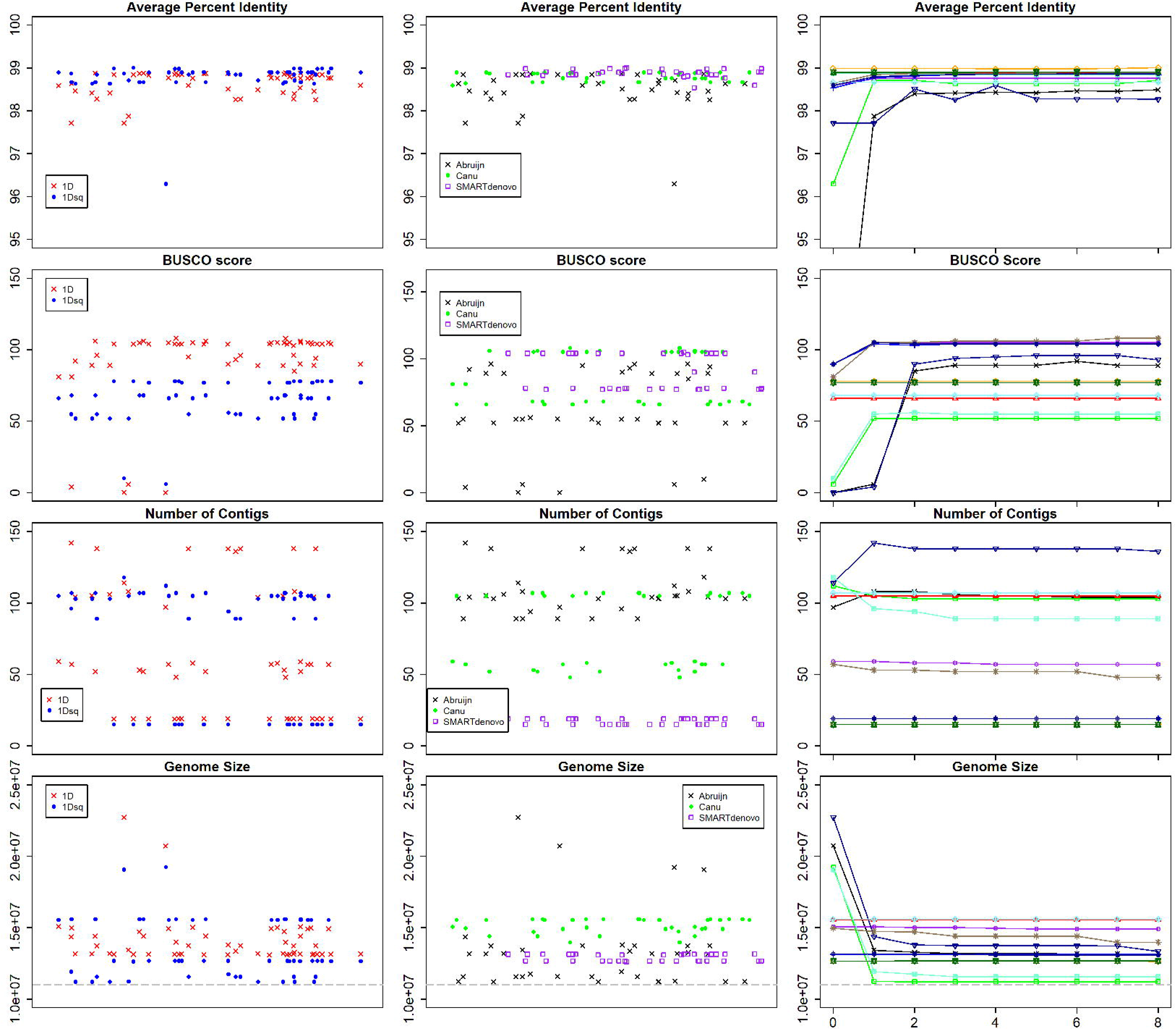
Performance metrics for all *Giardia* BGS long read assemblies. The title above each scatterplot denotes the metric being plotted on the y-axis. The left column shows the differences between 1D (red Xs) vs 1Dsq (blue circles) data for each assembly protocol. Note that the data are paired. The middle column shows the assemblies separated by assembly program: abruijn (black Xs), canu (green circles), and SMARTdenovo (purple boxes). In the left and middle columns, the assemblies are randomly assigned along the x-axis for visualization purposes, hence there are no units. The right column shows polished sets of assemblies with the x-axis denoting how many times the draft assembly was polished. The dashed grey line shows the size of the *Giardia* BGS reference assembly.

Using NanoOK [15], 1D reads were aligned to the corresponding reference genome and the error profiles of aligned reads were evaluated. NanoOK outputs read error profiles for each reference contig. To get overall error profiles for all reads, the values for each contig were multiplied by the proportion of total reads that aligned to that contig. The sum of these values for each error metric were scaled according to the proportion of total sequencing reads that were used for NanoOK’s analysis.

#### Hybrid assemblies

Hybrid assemblies for *Giardia* AWB were created from every AWB long read assembly in Supplementary Table 1. All of the AWB hybrid assemblies with the highest complete BUSCO score (117, Supplementary Table S11) were constructed from a SMARTdenovo long read assembly. For this reason, and because of the performance of the long read SMARTdenovo assemblies in general (See Discussion of long read assemblies), the *Giardia* BGS and beaver hybrid assemblies were constructed from Illumina reads and the SMARTdenovo assemblies of the 1D MinION reads. The AWB hybrid assemblies outperformed their long read counterparts in all metrics measured (Supplementary Tables S1 and S11) and, for all three isolates, the hybrid assemblies had higher complete BUSCO scores than their corresponding long read assembly. The best hybrid assembly for each isolate was selected for all further analysis on the basis of maximum complete BUSCO score (AWB_hybrid_106_0150015723312338_1dsmartx0, BGS_hybrid_gs3-20-2019_22372244_1dsmartx0, Beaver_hybrid_107218_2309_1dsmartx0). For each of these assemblies, alignment to the AWB reference genome showed that the full chromosome was recovered for chromosomes 1 – 4 and the majority of chromosome 5 was also recovered (Fig. 2).

**Figure 2.**
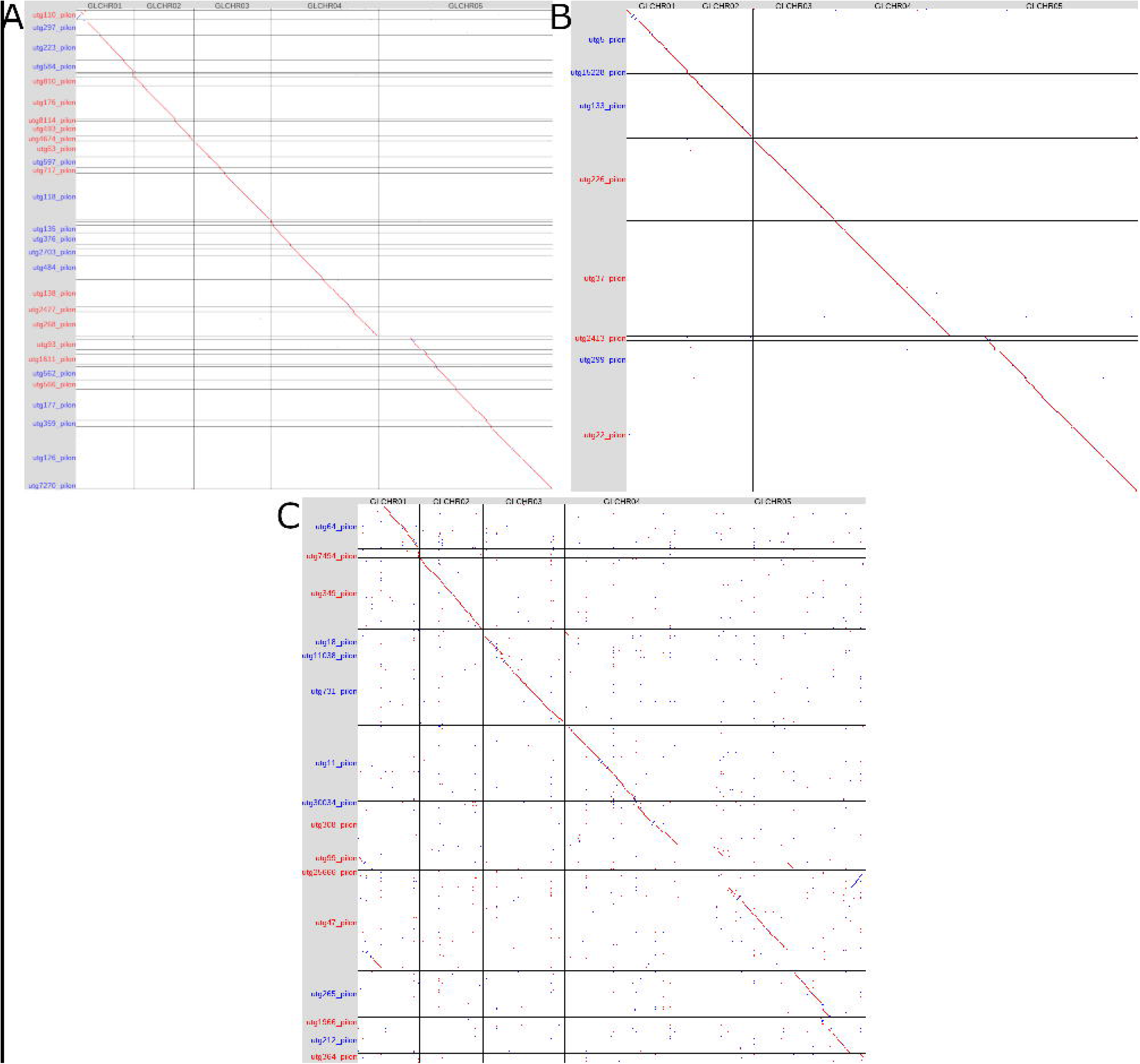
Dotplots (Oxford Grids) of pairwise whole genome alignments between the *Giardia* AWB reference genome and A) the *Giardia* AWB hybrid genome, B) the *Giardia* beaver hybrid genome, and C) the *Giardia* BGS hybrid genome. Each of the five *Giardia* chromosomes from the reference genome is represented as a column and each contig from the hybrid genome is represented as a row. Contig names and dots in the plot coloured red represent forward alignments while contig names and dots coloured in blue are reverse alignments.

### Structural variant analysis

We predicted structural variants from the long reads and hybrid assemblies to examine the variation between the four copies of each chromosome in the *Giardia* isolates sequenced. *Giardia* AWB, BGS, and beaver had 392, 1860, and 483 variants respectively (Table 3), which affect 2072, 4151, and 3423 genes respectively. For each isolate, the full lists of predicted structural variants and genes affected by each variant can be found in Supplementary Tables S12 – S14. Notably among the genes affected are known virulence factors including variant-specific surface proteins (VSP), tenascins, and high cysteine membrane proteins [16]. In AWB, BGS, and beaver 39, 97, and 56 of the structural variants were found to have significantly more VSP than expected, respectively. Figure 3 shows alignments of the three hybrid genomes to the AWB reference genome with the predicted structural variants for each genome.

**Table 3.** Structural variants (SVs) in *Giardia* AWB, BGS, and beaver. Numbers in brackets are average lengths (bp) of the variants.

|  | **AWB** | **BGS** | **beaver** |
| --- | --- | --- | --- |
| **Number of SVs** | 392 | 1860 | 483 |
| **# Duplications** | 45 (14520.4) | 185 (48239.6) | 69 (37535.0) |
| **# Deletions** | 46 (15487.1) | 298 (34454.6) | 74 (46361.1) |
| **# Inversions** | 162 (19437.9) | 746 (28782.2) | 234 (12866.7) |
| **# Inverted Duplications** | 2 (2257.0) | 14 (2680.1) | 0 (0.0) |
| **# Transversions** | 104 (2.3) | 436 (20.8) | 46 (4.0) |
| **# Insertions** | 33 (299.6) | 181 (596.4) | 60 (286.9) |
| **Proportion of genome contained in SVs** | 0.1876 | 0.5662 | 0.3372 |
| **Number of genes in SVs** | 2072 | 4151 | 3423 |

**Figure 3.**
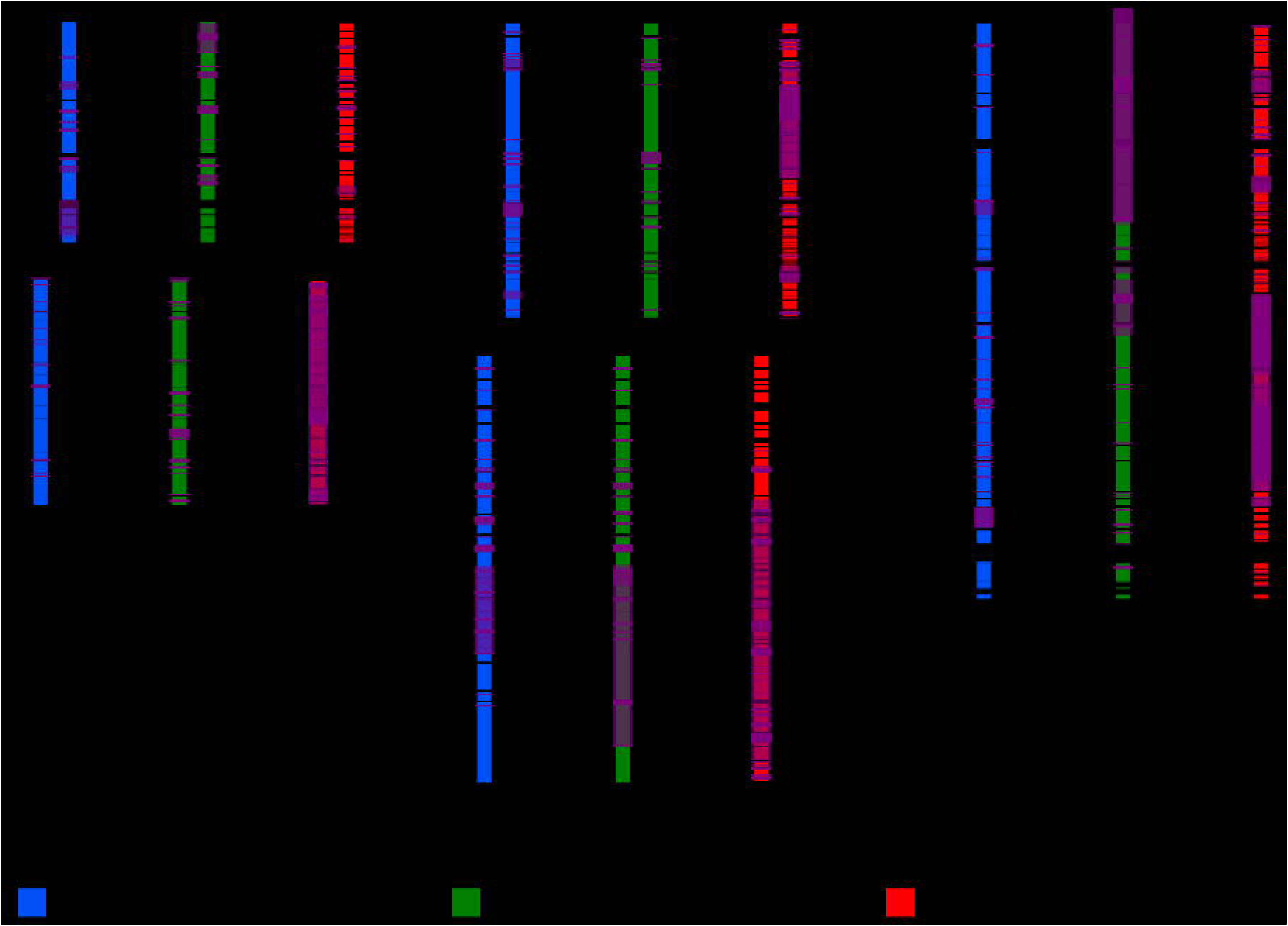
Whole genome alignments with predicted structural variants. The hybrid assembly contigs are shown as coloured boxes next to the reference *Giardia* AWB chromosome to which they align (black lines with vertical names beside each). Translucent purple boxes above the contigs show the locations and sizes of predicted structural variants in all three hybrid genomes. Note to reviewers: an interactive version of figure 3 that has filtering capabilities for viewing the structural variants can be found at: http://pages.cpsc.ucalgary.ca/~stephen.pollo/Giardia_SV_Fig/ This version would be added to the GigaScience GigaDB database linked to the paper

### *Genome of* Giardia *beaver*

The genome of *Giardia* beaver was assembled into 8 contigs totalling 11,467,485 bp. It has a maximum contig length of 2.759 Mb and an N50 of 1.965 Mb. One hundred thirteen complete BUSCOs were found out of 134 detected across the three *Giardia* isolates examined here. *Giardia* beaver has 49.56% GC content, similar to values found for *Giardia* AWB (49.0) and other assemblage A isolates (49.25; 49.04) [2,17].

## Discussion

### Long read assemblies and assemblers that lead to reference quality hybrid assemblies

Among the three assemblers tested, the SMARTdenovo assemblies for both *Giardia* AWB and BGS showed the lowest variability in all metrics except average indel size (Fig. 1 and Supplementary Figs. S1 – S10). Moreover, the SMARTdenovo assemblies had the highest average values for average percent identity, BUSCO score, and proportion of reference covered 1X (where higher values indicate better performance) (Supplementary Table S1) and consistently strong performance in all metrics except average indel size (Fig. 1 and Supplementary Figs. S1 - S10). Despite thirteen of the top performing assemblies (8 AWB, 5 BGS) being Abruijn assemblies (Supplementary Table S2), plotting values for each metric showed Abruijn had the most variable performance (Supplementary Figs. S1 – S10, Supplementary Tables S7 – S8). Canu assemblies generally performed somewhere between the SMARTdenovo and Abruijn assemblies (Supplementary Tables S7 – S8).

Analysis of the 207 AWB and 108 BGS assemblies indicates that the optimal long read only assembly pipeline for MinION sequenced *Giardia* is a SMARTdenovo assembly from 1D reads (either pooled or non-pooled input to reach sufficient genome coverage) followed by four or five rounds of polishing with Nanopolish (See Supplementary Material for discussion of 1D vs 1Dsq input reads, pooling different sequencing runs for the same organism, and number of rounds of genome polishing). However, it was the unpolished long read assemblies that resulted in the best hybrid assemblies (1D read, SMARTdenovo assembled, no polishing with Nanopolish; Supplementary Table S11). Interestingly, the BGS assemblies are larger than the reference BGS assembly that was generated from 454 data [4], potentially due to the fragmented nature of the reference assembly. The AWB and BGS hybrid assemblies generated here have higher complete BUSCO scores than the available reference genomes (117 for both hybrids vs 114 AWB reference and 116 BGS reference) and were assembled into very large pieces (AWB hybrid N50: 616 kb; BGS hybrid N50: 1,645 kb), suggesting they are of reference quality (Figs. 2 and 3). Moreover, the hybrid genome for *Giardia* beaver has a similarly high complete BUSCO score and similar contig numbers and contig lengths to the AWB and BGS hybrids, indicating that reference quality assemblies can be generated *de novo* for *Giardia* with as little as one ONT MinION and one multiplexed Illumina MiSeq sequencing run.

An optimal assembly pipeline for MinION data can change with each release of new programs specializing in handling long error prone reads. Already having the scripts to calculate the evaluation metrics used here makes re-evaluations easier to perform and enables evaluation of assembler performance that is current with each new program or version release. The typical publication process, from numerous drafts of a manuscript and peer-review, can be time-consuming and not conducive to keeping such an analysis current. Therefore, a blog or community forum similar to an analysis on github of MinION basecalling programs [18] would be more appropriate. These media may also make it easier to discuss issues surrounding installation of these programs and running them in various computing environments. For example, some of the programs used here took up to a month to get installed and running properly. Having a current analysis of available long read assemblers would therefore also allow researchers to determine which programs are worth the time to get working and when it may be a better use of time to go with programs that need less configuration (like Canu which worked immediately) but will still perform adequately for the intended purpose.

### Structural variants reveal different levels of intra-isolate variation

Despite having similar genome sizes, the three isolates examined here have very different total numbers of variants detected and proportions of their genomes that are within a structural variant region (Table 3, Fig. 3). When *Giardia* BGS was first sequenced, the authors noted a much higher allelic sequence heterozygosity than what was observed in AWB (0.53% in BGS vs 0.01% in AWB) [4]. The same trend is observed in the structural variants here with BGS being considerably more heterozygous than AWB. The differences in allelic sequence heterozygosity were attributed to AWB and BGS being in different assemblages [4]. While the values for *Giardia* beaver (an assemblage A isolate) being more similar to AWB than BGS (Table 3) tentatively support the hypothesis that assemblage B is more heterozygous than assemblage A, many more genomes from each assemblage are needed to confirm it. Further, single cell sequencing could be used to examine the population structure of the isolates at a genetic level. Nonetheless, assemblage-specific variations in heterozygosity, or even isolate-specific variations in heterozygosity, will be important to consider in future comparisons between *Giardia* genomes. Previous genomic comparisons between assemblages [4] and within assemblages [19] have focused on SNPs and analyses of specific gene families. Including structural variant information provides a more complete picture of the heterozygosity and genetic diversity of each isolate by capturing differences in gene dosage as well as gene content.

### Effects of recombination in *Giardia* on structural variants

Recombination between different cells (outcrossing) within and between isolates of *Giardia* has been suggested to occur through an as-yet undiscovered mechanism [20–23]. Outcrossing recombination events would allow for changes in gene copy number if the event involved or encompassed a structural variant like a duplication or deletion. Alternatively, large inversions can prevent recombination in the inverted areas [24], preventing gene flow during recombination events in *Giardia*. These regions are therefore important to keep in mind in future studies on recombination in *Giardia* as they may confound the analyses. Several dozen structural variants from each of the isolates examined here were found to be significantly enriched for VSP, supporting the suggestion that recombination is a potential source of VSP variation [25]. Expansions and contractions of this gene family through inheritance during outcrossing events of duplicated or deleted loci that affect VSP could be an important factor in the number and distribution of these genes between the various *Giardia* assemblages and isolates. As key surface proteins involved in host immune evasion [26], these expansions and contractions of the VSP repertoire could partially explain differences in pathogenicity between isolates. Moreover, as mediators of the *Giardia* cell’s interaction with its surrounding environment, expansions and contractions of the VSP repertoire could affect host range. Alternatively, these genes could be hotspots for recombination events that generate structural variants. Then in addition to their roles as surface proteins they would also be potential factors influencing the evolution of *Giardia* genomes.

## Conclusions

The present study demonstrates that high quality genomes can be generated for *Giardia* for a few thousand dollars per genome, thus enabling future large scale comparative genomic studies of the genus. Moreover, third generation long reads can be further used to investigate heterozygosity and genome organization in *Giardia* despite its tetraploidy. We showed that structural variant regions affect many genes notably virulence factors including VSP, suggesting an important mechanism in the inheritance and distribution of these proteins among *Giardia* isolates. Finally, we have generated a reference genome sequence for a new isolate, *Giardia* beaver, with accompanying prediction of its structural variants.

## Methods

### Giardia duodenalis isolates

*Giardia* AWB (ATCC 30957) and *Giardia* BGS (ATCC 50580) were obtained from the American Tissue Culture Collection, while *Giardia* beaver was a gift from Dr. Gaetan Faubert from McGill University. *Giardia* trophozoites were grown in TYI-S-33 medium [27] in 16-mL screw capped glass tubes incubated at 37°C.

### DNA extraction

Ten 16-mL culture tubes of each *Giardia* isolate (AWB, BGS, and beaver) grown to late logarithm stage (~5 - 8 × 10^5 cells/mL) were used for genomic DNA isolation. The culture tubes were chilled on ice for 5 min and the cells were collected by centrifugation at 1,100 × g for 15 min at 4°C. Genomic DNA was extracted with DNAzol Reagent (ThermoFisher Scientific) by following the manufacturer’s instructions. Briefly, each cell pellet was resuspended and lysed in DNAzol Reagent by gentle pipetting followed by a freeze (30 min at 80°C) and thaw (10 min at room temperature) step. The lysate was then centrifuged at 10,000 × g for 10 min at 4°C to remove insoluble cell debris. The supernatant was transferred to a new tube and the DNA was recovered by centrifugation of the supernatant at 4,000 × g for 5 min at 4°C. The DNA pellet was washed twice with 75% ethanol then air-dried. The DNA was resuspended initially in 8 mM NaOH then neutralized by addition of HEPES to a final concentration of 9 mM.

RNA was removed from the DNA sample by the addition of 1 - 2 μL of 20 μg/μL RNase A (BioShop) followed by incubation at 65°C for 10 min. The degraded RNA was precipitated by the addition of ammonium acetate, incubation at 4°C for 20 min, and centrifugation at 12,000 × g for 30 min at 4°C. The supernatant was transferred to a new tube and the DNA was precipitated by the addition of 95% ethanol, incubation at room temperature for 5 min, and centrifugation at 12,000 × g for 20 min at 4° C. The DNA pellet was washed once with 0.01M ammonium acetate in 75% ethanol and once with 75% ethanol alone. The DNA pellet was air-dried before resuspension in TE buffer (10 mM Tris-HCl pH 8.0, 1 mM EDTA).

### MinION sequencing

The 1Dsq library preparation kit SQK-LSK308 was used as recommended by the manufacturer (Oxford Nanopore Technologies, Oxford, United Kingdom). Approximately 200 ng of prepared library was loaded onto a FLO-MIN107 (R9.5) flow cell. Data collection was carried out with live basecalling for 48 h, or until no more strands were being sequenced. All sequences were deposited in the sequence read archive (SRA) under accession number PRJNA561185.

### Illumina sequencing

Libraries were prepared using NexteraXT and paired-end sequenced on the MiSeq (v3, 2×300 cycles) or iSeq 100 (I1, 2×150 cycles) platforms according to manufacturer instructions (Illumina Inc). All sequences were deposited in the SRA under accession number PRJNA561185.

### Long read basecalling, *de novo* assembly, and genome polishing

Basecalling of all MinION output files was performed with the program Albacore (version 2.0.2) [28] using the full_1dsq_basecaller.py method to basecall both 1D and 1Dsq reads. The flowcell and kit parameters were FLO-MIN107 and SQK-LSK308 respectively. The general command used to run Albacore was: `full_1dsq_basecaller.py --flowcell FLO-MIN107 --kit SQK-LSK308 --input PATH/TO/FAST5/FILES --save_path ./ --worker_threads 38`

*De novo* assemblies were performed using the programs Abruijn (version 2.1b) [29], Canu (version 1.6) [30], and SMARTdenovo (version 1.11 running under Perl version 5.22.0) [31]. Abruijn assemblies were conducted using the nanopore platform setting, coverage estimates calculated as the number of bases in the input reads divided by the reference genome size (Table 1) all rounded to the nearest integer, and all other default settings (one polishing iteration, automatic detection of kmer size, minimum required overlap between reads of 5000 bp, automatic detection of minimum required kmer coverage, automatic detection of maximum allowed kmer coverage). Canu assemblies were performed using Canu’s settings for uncorrected nanopore reads (-nanopore-raw), genome sizes estimated from the reference genome sizes (Table 1), and setting gnuplotTested=true to bypass html output report construction. SMARTdenovo assemblies were conducted using default settings (kmer length for overlapping of 16 and minimum required read length of 5000 bases). The general commands used to run each of the assemblers, with variable parameters written in upper case, were:

Abruijn: `abruijn PATH/TO/READS out_nano COVERAGE_ESTIMATE -- platform nano --threads 56`
Canu: `canu -p UNIQUE_NAME genomeSize=12.8m -nanopore-raw PATH/TO/READS gnuplotTested=true`
SMARTdenovo: `smartdenovo.pl -p UNIQUE_NAME PATH/TO/READS > UNIQUE_NAME.mak, followed by the command: make -f UNIQUE_NAME.mak`

Genome polishing is an error correction step performed on assemblies generated from third-generation data to compensate for the high error rate of the reads [8]. It involves re-evaluating the base calls from the MinION squiggle files together with the read overlap information from the assembly to improve base accuracy and correct small insertions and deletions [32]. Here polishing was performed with the program Nanopolish (version 0.8.5) following the directions for “computing a new consensus sequence for a draft assembly” [33]. Briefly, the draft genome was first indexed using BWA (version 0.7.15-r1140) [34] and the basecalled reads were aligned to the draft genome using BWA. SAMtools (version 1.6 using htslib 1.6) [35] was then used to sort and index the alignment. Nanopolish then computed the new consensus sequence in 50kb blocks in parallel, which were then merged into the polished assembly. The general commands used to run Nanopolish were:

~~~
nanopolish index -d PATH/TO/FAST5/FILES PATH/TO/READS
bwa index PATH/TO/ASSEMBLY/TO/POLISH
bwa mem -x ont2d -t 8 PATH/TO/ASSEMBLY/TO/POLISH PATH/TO/READS |
samtools sort -o reads.sorted.bam -T reads.tmp
samtools index reads.sorted.bam
python ~/nanopolish/scripts/nanopolish_makerange.py
PATH/TO/ASSEMBLY/TO/POLISH | parallel --results
nanopolish.results -P 14 nanopolish variants --consensus
UNIQUE_NAME_polished_x${POLISHING_ITERATION}.{1}.fa -w {1} -r
PATH/TO/READS -b reads.sorted.bam -g PATH/TO/ASSEMBLY/TO/POLISH
-t 4 --min-candidate-frequency 0.1
python ~/nanopolish/scripts/nanopolish_merge.py
UNIQUE_NAME_polished_x${POLISHING_ITERATION}.*.fa >
UNIQUE_NAME_polished_x${POLISHING_ITERATION}_genome.fa
~~~

### Read error profile analysis

Read error profiles were examined for the six *Giardia* AWB and *Giardia* BGS runs using the program NanoOK (version v1.31) [15]. NanoOK extracts fasta sequences from the fast5 files produced by the MinION and aligns them to the reference genome using the LAST aligner (version 876) [36]. It then calculates error profiles for each set of reads that aligned to each contig in the reference. To obtain overall values for all reads in the sequencing run, for each error metric the value for each contig was extracted from the .tex file produced by NanoOK and multiplied by the proportion of the total reads mapping to that contig. These values were then summed to yield the metric value with respect to all reads in the sequencing run. The sums were scaled according to the proportion of the total reads that were included in the metric calculation - those that were mapped to the contigs - to yield the metric value for all reads used in the analysis.

### Long read assembly evaluation

The effects on final assembly quality were evaluated for the following parameters: 1D vs 1Dsq input reads, pooling reads for the same organism from multiple runs, assembly program, and number of genome polishing iterations. Firstly, 13 distinct input combinations, that represent all permutations of pooling runs for the same organism for both 1D and 1Dsq reads, were used for *de novo* assemblies: AWB_0157 1D reads, AWB_0157 1Dsq reads, AWB_0150_0157 1D reads, AWB_0150_0157 1Dsq reads, AWB_2338 1D reads, AWB_2338 1Dsq reads, AWB_2331_2338 1D reads, AWB_0150_0157_2331_2338 1D reads, AWB_0150_0157_2338 1Dsq reads, BGS_2244 1D reads, BGS_2244 1Dsq reads, BGS_2237_2244 1D reads, and BGS_2237_2244 1Dsq reads (Table 1). Each of these input combinations was used to perform a *de novo* assembly with each of the three assemblers used: Abruijn, Canu, and SMARTdenovo. All of the resulting assemblies that produced contiguous sequences were polished with Nanopolish. Eight rounds of Nanopolish polishing were performed on the Canu and SMARTdenovo assemblies and seven rounds were performed on the Abruijn assemblies (which get polished once by Abruijn).

All assemblies and polished versions of the assemblies were aligned to the corresponding reference genome using the LAST aligner (version 876) [36] following the example for human-ape alignments [37]. Briefly, the reference genome was indexed using LAST, then substitution and gap frequencies were determined using the last-train method [38]. Finally, alignments were performed using the lastal method and the determined substitution and gap frequencies. The resulting alignments were then filtered to retain only those alignments with an error probability < 1e^−5^. *Giardia* AWB assemblies were aligned to only the contigs from the reference genome labelled GLCHR01, GLCHR02, GLCHR03, GLCHR04, and GLCHR05 (representing the five chromosomes of *Giardia duodenalis*). Filtered alignments were converted to other file formats (for metric calculation) using the maf-convert method in the LAST aligner.

Average percent identity was calculated from alignments in blasttab format by taking the sum of the percent identity multiplied by the alignment length for each aligned portion and dividing that sum by the total alignment length. Proportion of mismatching bases was calculated from alignments in psl format by taking the sum of mismatching bases for all aligned portions divided by the total alignment length. Total number of indels per 1000 aligned bases was calculated from alignments in psl format by taking the sum of the number of insertions in the query and the number of insertions in the target for all aligned portions, dividing that sum by the total alignment length and multiplying by 1000. Average size of indels was calculated from alignments in psl format by taking the sum of the number of bases inserted in the query and the number of bases inserted in the target for all aligned portions and dividing that sum by the total number of indels. The proportions of the reference covered 0, 1, 2, 3, or 4 times were calculated using BEDtools (version v2.27.1) [39]. Alignments were first converted to SAM format and SAMtools was used to sort the alignment and convert it to a bam file. The genomecov function of BEDtools was then used to analyze the coverage of every base in the reference genome in the alignment. The proportion of bases in the reference genome with 0, 1, 2, 3, and 4 fold coverage in the assembly were retrieved.

The assembly evaluation metrics Number of Contigs and Genome Size were calculated for each assembly from the assembly fasta file. BUSCOs were calculated for each assembly using BUSCO v3.0.2 (BLAST+ v2.6.0, HMMER v3.1b2, and AUGUSTUS v3.2.3), with the eukaryote_odb9 dataset and default options (-sp fly) [40].

Average and standard deviation values for the groupings presented in the tables and figures for each metric were calculated in R [41]. R was also used to construct the scatter plots for the figures.

### Hybrid assemblies

Hybrid genome assemblies were generated using the program Pilon (version 1.22) [42]. Briefly, short, highly accurate reads are mapped to a long-read assembly to correct for the higher error rate in the long reads. For each hybrid assembly, the Illumina reads were mapped using BWA to the long read assembly. After sorting and indexing the alignments with SAMtools, pilon was run with default parameters to generate the hybrid assemblies. The general command to run pilon was:

~~~
pilon -Xm×200g --genome GENOME_TO_CORRECT --frags
BAM1.sorted.bam --frags BAM2.sorted.bam --output UNIQUE_NAME
~~~

The improvement of the hybrid assembly over the long read assembly from which it was built was measured by the BUSCO scores of each (calculated as described above). BUSCO scores were preferred because they do not depend on having a reference sequence and gene finding depends on assembly accuracy. The best hybrid assembly for each isolate was deposited at DDBJ/ENA/GenBank under the accession numbers VSRS00000000 (*Giardia* beaver), VSRT00000000 (*Giardia* AWB), and VSRU00000000 (*Giardia* BGS). The versions described in this paper are versions VSRS01000000, VSRT01000000, and VSRU01000000 respectively.

### Structural variant prediction and analysis

Structural variants were predicted using the programs ngmlr and sniffles [10]. For each *Giardia* isolate, the long reads were mapped to the best hybrid assembly using ngmlr v0.2.7. The resulting alignments were sorted with SAMtools and the variants were called with sniffles v1.0.10. The general commands to run ngmlr and sniffles were:

~~~
ngmlr -t 56 -r HYBRID_ASSEMBLY -q LONG_READS -o
UNIQUE_NAME_ngmlr.sam -x ont
sniffles -t 56 --genotype --cluster --report_seq -n -1 -m
ALIGNED_LONG_READS_ngmlr_sorted.bam -v UNIQUE_NAME_SVs.vcf
~~~

Genes likely to be affected by the structural variants were identified by mapping known proteins from the *Giardia* AWB reference genome to the hybrid assembly used to predict the structural variants with the program exonerate v2.2.0 [43] and finding the genes overlapping the variant regions using BEDtools. The general commands were:

~~~
exonerate -m protein2genome -q AWB_PROTEINS.gff -t
HYBRID_ASSEMBLY.fasta -M 250000 -n 1 --showalignment FALSE --
showvulgar FALSE --showtargetgff > UNIQUE_NAME.txt
sed ‘/^#/d’ UNIQUE_NAME.txt > UNIQUE_NAME.gff
sed ‘1,2d;$d’ UNIQUE_NAME.gff > UNIQUE_NAME_2.gff
bedtools intersect -a UNIQUE_NAME_SVs.vcf -b UNIQUE_NAME_2.gff -
wb > UNIQUE_NAME_intersect_vcf_genesonlyn1gff.txt
~~~

For each variant type, the list of putatively affected genes was examined and genes of interest were analyzed for enrichment in the variants. For each predicted variant, 10000 random samples of the same size as the variant were selected from the genome. For each sample the overlapping genes were found and the genes of interest were counted. The 95^th^ percentile was calculated from the resulting distribution of genes of interest using the nearest-rank method to find the count above which there is significant enrichment of the gene of interest (ie. the cutoff for rejecting H_0_). The subsampling experiment was implemented in Java, the code for which is available on github at https://github.com/StephenMJPollo/SV_Subsampling.

### Genome assembly for *Giardia* beaver

The genome of *Giardia* beaver was assembled *de novo* from 1D minION reads using SMARTdenovo (see discussion; commands are the same as in methods above). Illumina reads were added to create a hybrid assembly as described above.

## Availability of source code and requirements

Project name: SV_Subsampling

Project home page: https://github.com/StephenMJPollo/SV_Subsampling

Operating system: Linux

Programming Language: Java

Other requirements: BEDtools

## Availability of supporting data and materials

Sequence reads are available on the SRA under accession number PRJNA561185. The hybrid assemblies generated are available from GenBank under the accession numbers VSRS00000000 (*Giardia* beaver), VSRT00000000 (*Giardia* AWB), and VSRU00000000 (*Giardia* BGS). The versions described in this paper are versions VSRS01000000, VSRT01000000, and VSRU01000000 respectively. All other supporting material will be submitted to the GigaScience GigaDB database.

## Additional Files

Supplementary_Discussion: Additional discussion on long read only assemblies.

Supplementary Figures: Figures S1 – S10 with corresponding legends.

Supplementary Tables: Tables S1 – S15.

## List of abbreviations

bp: base pairs
BUSCO: benchmarking universal single copy orthologs
ONT: Oxford Nanopore Technologies
SNPs: single nucleotide polymorphisms
SRA: sequence read archive
SVs: structural variants
VSP: variant-specific surface proteins

## Consent for publication

Not applicable.

## Authors’ contributions

Resources and investigation: GWB, JW, and SJR. Investigation, formal analysis, software, writing original draft, and visualization: SMJP. Funding acquisition and supervision: RAG, JY, BRD, and JDW. Conceptualization and methodology: SMJP, SJR, MLW, RAG, BRD, and JDW.

## Acknowledgements

Not applicable

## References

1. Certad G, Viscogliosi E, Chabé M, Cacciò SM. Pathogenic mechanisms of *Cryptosporidium* and *Giardia*. Trends Parasitol. 2017;33:561–76.

2. Morrison HG, McArthur AG, Gillin FD, Aley SB, Adam RD, Olsen GJ, et al. Genomic minimalism in the early diverging intestinal parasite *Giardia lamblia*. Science. 2007;317:1921–6.

3. Aurrecoechea C, Brestelli J, Brunk BP, Carlton JM, Dommer J, Fischer S, et al. GiardiaDB and TrichDB: Integrated genomic resources for the eukaryotic protist pathogens *Giardia lamblia* and *Trichomonas vaginalis*. Nucleic Acids Res. 2009;37:526–30.

4. Franzén O, Jerlström-Hultqvist J, Castro E, Sherwood E, Ankarklev J, Reiner DS, et al. Draft genome sequencing of *Giardia intestinalis* Assemblage B isolate GS: Is human giardiasis caused by two different species? Plos Pathog. 2009;5:e1000560.

5. Stroehlein AJ, Korhonen PK, Chong TM, Lim YL, Chan KG, Webster B, et al. High-quality *Schistosoma haematobium* genome achieved by single-molecule and long-range sequencing. Gigascience. 2019;8:1–12.

6. Rhoads A, Au KF. PacBio sequencing and its applications. Genomics Proteomics Bioinformatics. 2015;13:278–89.

7. Tyson JR, O’Neil NJ, Jain M, Olsen HE, Hieter P, Snutch TP. Whole genome sequencing and assembly of a *Caenorhabditis elegans* genome with complex genomic rearrangements using the MinION sequencing device. bioRxiv. 2017;

8. Lu H, Giordano F, Ning Z. Oxford nanopore minION sequencing and genome assembly. Genomics Proteomics Bioinformatics. 2016;14:265–79.

9. de Koning APJ, Gu W, Castoe TA, Batzer MA, Pollock DD. Repetitive elements may comprise over two-thirds of the human genome. PLoS Genet. 2011;7:e1002384.

10. Sedlazeck FJ, Rescheneder P, Smolka M, Fang H, Nattestad M, Von Haeseler A, et al. Accurate detection of complex structural variations using single-molecule sequencing. Nat Methods. 2018;15:461–8.

11. Jeffares DC, Jolly C, Hoti M, Speed D, Shaw L, Rallis C, et al. Transient structural variations have strong effects on quantitative traits and reproductive isolation in fission yeast. Nat Commun. 2017;8:14061.

12. Weischenfeldt J, Symmons O, Spitz F, Korbel JO. Phenotypic impact of genomic structural variation: insights from and for human disease. Nat Rev Genet. 2013;14:125–38.

13. Kyriakidou M, Tai HH, Anglin NL, Ellis D, Strömvik M V. Current Strategies of Polyploid Plant Genome Sequence Assembly. Front Plant Sci. 2018;9:1–15.

14. Feng Y, Zhang Y, Ying C, Wang D, Du C. Nanopore-based fourth-generation DNA sequencing technology. Genomics Proteomics Bioinformatics. 2015;13:4–16.

15. Leggett RM, Heavens D, Caccamo M, Clark MD, Davey RP. NanoOK: Multi-reference alignment analysis of nanopore sequencing data, quality and error profiles. Bioinformatics. 2016;32:142–4.

16. Dubourg A, Xia D, Winpenny JP, Naimi S Al, Bouzid M, Sexton DW, et al. *Giardia* secretome highlights secreted tenascins as a key component of pathogenesis. Gigascience. 2018;7:1–13.

17. Adam RD, Dahlstrom EW, Martens CA, Bruno DP, Barbian KD, Ricklefs SM, et al. Genome Sequencing of *Giardia lamblia* Genotypes A2 and B Isolates (DH and GS) and Comparative Analysis with the Genomes of Genotypes A1 and E (WB and Pig). Genome Biol Evol. 2013;5:2498–511.

18. Wick R. A comparison of different Oxford Nanopore basecallers. 2017. https://github.com/rrwick/Basecalling-comparison#m.

19. Ankarklev J, Franzén O, Peirasmaki D, Jerlström-Hultqvist J, Lebbad M, Andersson J, et al. Comparative genomic analyses of freshly isolated *Giardia intestinalis* assemblage A isolates. BMC Genomics. 2015;16:1–14.

20. Cooper MA, Sterling CR, Gilman RH, Cama V, Ortega Y, Adam RD. Molecular Analysis of Household Transmission of *Giardia lamblia* in a Region of High Endemicity in Peru. J Infect Dis. 2010;202:1713–21.

21. Cooper MA, Adam RD, Worobey M, Sterling CR. Population Genetics Provides Evidence for Recombination in *Giardia*. Curr Biol. 2007;17:1984–8.

22. Ankarklev J, Lebbad M, Einarsson E, Franzén O, Ahola H, Troell K, et al. A novel high-resolution multilocus sequence typing of *Giardia intestinalis* Assemblage A isolates reveals zoonotic transmission, clonal outbreaks and recombination. Infect Genet Evol. 2018;60:7–16.

23. Birky CW. *Giardia* Sex? Yes, but how and how much? Trends Parasitol. 2010. p. 70–4.

24. Wellenreuther M, Mérot C, Berdan E, Bernatchez L. Going beyond SNPs: The role of structural genomic variants in adaptive evolution and species diversification. Mol Ecol. 2019;28:1203–9.

25. Jerlström-hultqvist J, Franzén O, Ankarklev J, Xu F, Nohýnková E, Andersson JO, et al. Genome analysis and comparative genomics of a *Giardia intestinalis* assemblage E isolate. BMC Genomics. 2010;11:543–58.

26. Prucca CG, Slavin I, Quiroga R, Elías E V., Rivero FD, Saura A, et al. Antigenic variation in *Giardia lamblia* is regulated by RNA interference. Nature. 2008;456:750–4.

27. Clark CG, Diamond LS. Methods for Cultivation of Luminal Parasitic Protists of Clinical Importance. Clin mi. 2002;15:329–41.

28. Vera D. Dockerfile for the Albacore basecaller from Oxford Nanopore. 2017. https://github.com/dvera/albacore.

29. Lin Y, Yuan J, Kolmogorov M, Shen MW, Chaisson M, Pevzner PA. Assembly of long error-prone reads using de Bruijn graphs. Proc Natl Acad Sci. 2016;113:E8396–405.

30. Koren S, Walenz BP, Berlin K, Miller JR, Bergman NH, Phillippy AM. Canu: scalable and accurate long-read assembly via adaptive *k*-mer weighting and repeat separation. Genome Res. 2017;27:722–36.

31. Ruan J. Ultra-fast de novo assembler using long noisy reads. 2017. https://github.com/ruanjue/smartdenovo.

32. Loman NJ, Quick J, Simpson JT. A complete bacterial genome assembled *de novo* using only nanopore sequencing data. Nat Methods. 2015;12:733–6.

33. Simpson J. Signal-level algorithms for MinION data. 2017. https://github.com/jts/nanopolish.

34. Li H, Durbin R. Fast and accurate long-read alignment with Burrows-Wheeler transform. Bioinformatics. 2010;26:589–95.

35. Cock PJA, Bonfield JK, Chevreux B, Li H. SAM/BAM format v1.5 extensions for *de novo* assemblies. bioRxiv. 2015;00:1–3.

36. Kielbasa SM, Wan R, Sato K, Horton P, Frith MC. Adaptive seeds tame genomic sequence comparison. Genome Res. 2011;21:487–93.

37. Mcfrith. last-genome-alignments. 2017. https://github.com/mcfrith/last-genome-alignments.

38. Hamada M, Ono Y, Asai K, Frith MC. Training alignment parameters for arbitrary sequencers with LAST-TRAIN. Bioinformatics. 2017;33:926–8.

39. Quinlan AR, Hall IM. BEDTools: A flexible suite of utilities for comparing genomic features. Bioinformatics. 2010;26:841–2.

40. Simão FA, Waterhouse RM, Ioannidis P, Kriventseva E V, Zdobnov EM. BUSCO: Assessing genome assembly and annotation completeness with single-copy orthologs. Bioinformatics. 2015;31:3210–2.

41. R Core Team. R: A language and environment for statistical computing. 2013. http://www.r-project.org/.

42. Walker BJ, Abeel T, Shea T, Priest M, Abouelliel A, Sakthikumar S, et al. Pilon: An integrated tool for comprehensive microbial variant detection and genome assembly improvement. PLoS One. 2014;9:e112963.

43. Slater GSC, Birney E. Automated generation of heuristics for biological sequence comparison. BMC Bioinformatics. 2005;6:1–11.

