## Supplemental Figures for "Nanopore sequencing of *Giardia* reveals widespread intra-isolate structural variation"

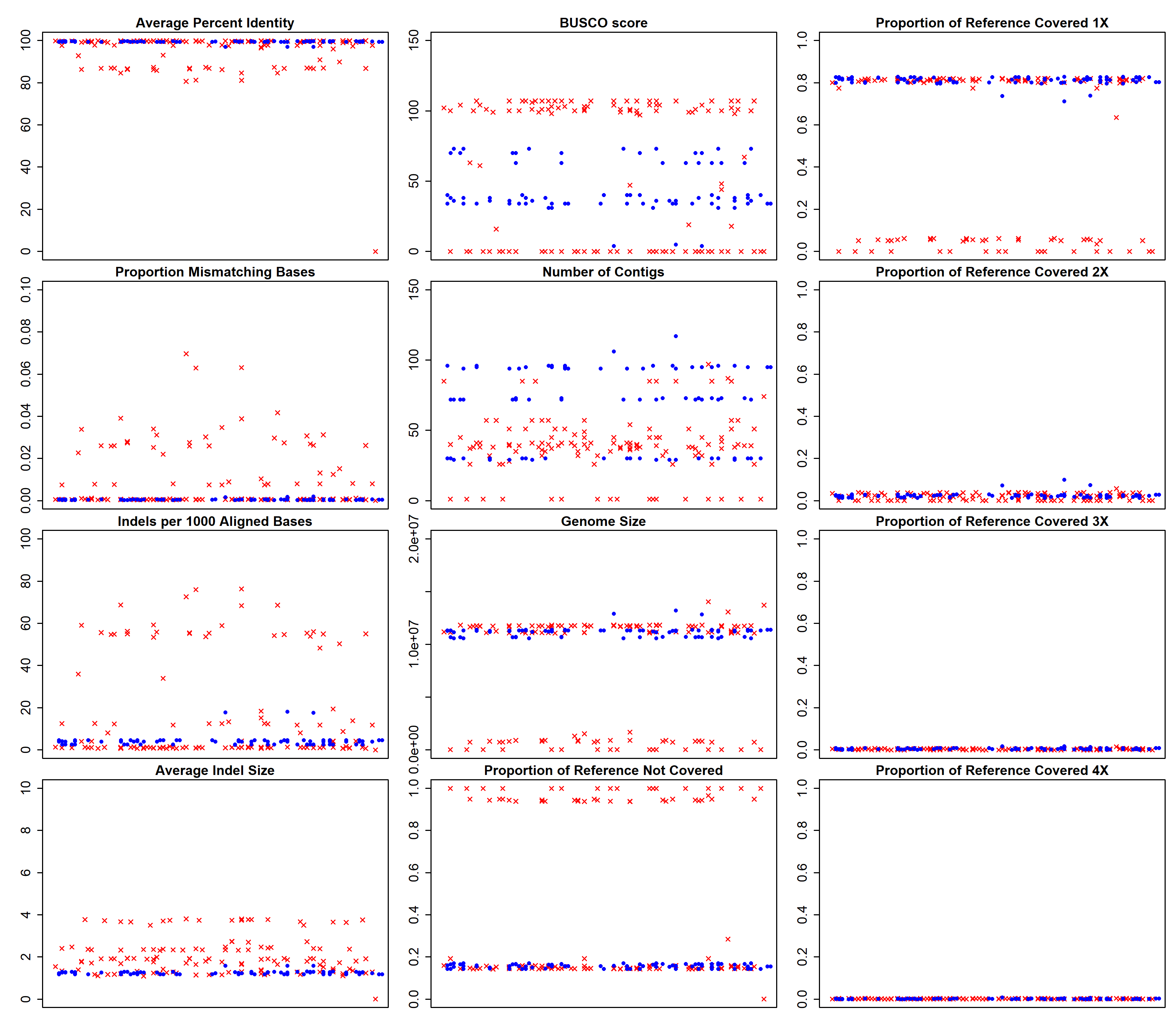
**Supplementary Figures**

**Figure S1.** Performance metrics for all 1D and 1Dsq *Giardia* AWB assemblies. Red X’s denote all assemblies created from 1D reads and blue circles denote all assemblies created from 1Dsq reads. The title above each scatterplot denotes the metric being plotted on the y-axis. The x-axis has no units because the x-values are randomly assigned to spread out the data points for visualization purposes. All metrics were calculated as described in the main text Methods. Assemblies from Run2 data are also included, even though their poor performance could be due to insufficient sequencing depth.

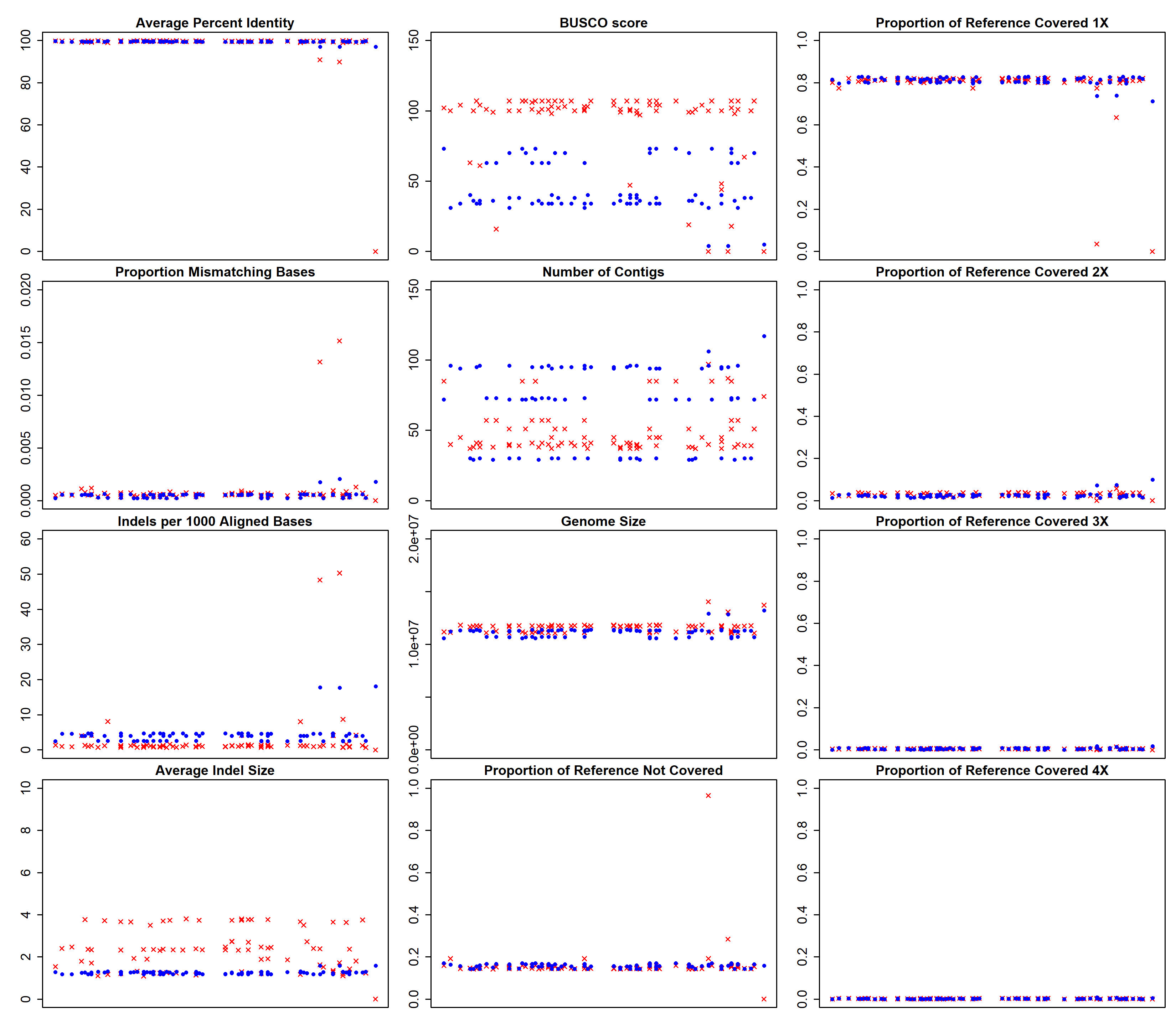

**Figure S2.** Performance metrics for corresponding 1D and 1Dsq *Giardia* AWB assembly pairs. All assemblies made from 1Dsq reads had a corresponding assembly made from 1D reads from the same inputs. Each pair was assigned the same random x-value so that corresponding 1D and 1Dsq assemblies would stack on top of each other in each plot. The additional assemblies from 1D reads are not shown. Red X’s denote all assemblies created from 1D reads and blue circles denote all assemblies created from 1Dsq reads. The title above each scatterplot denotes the metric being plotted on the y-axis. The x-axis has no units because the x-values are randomly assigned to spread out the data points for visualization purposes. All metrics were calculated as described in the main text Methods. Assemblies from Run2 data are also included, even though their poor performance could be due to insufficient sequencing depth.

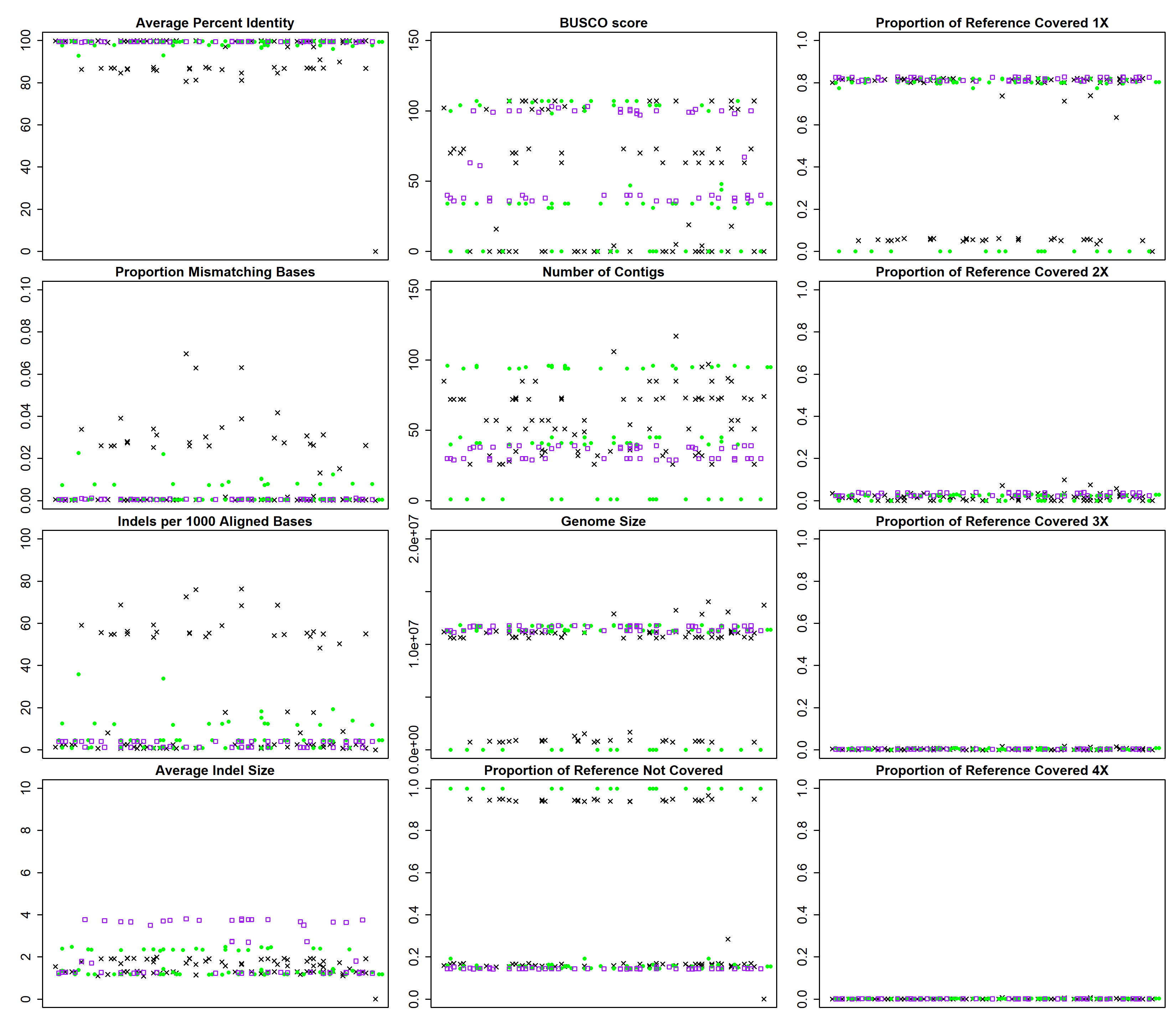

**Figure S3.** Performance metrics for all *Giardia* AWB assemblies, separated by assembly program. Black X’s denote all Abruijn assemblies, green circles denote all Canu assemblies, and purple squares denote all SMARTdenovo assemblies. The title above each scatterplot denotes the metric being plotted on the y-axis. The x-axis has no units because the x-values are randomly assigned to spread out the data points for visualization purposes. All metrics were calculated as described in the main text Methods. Assemblies from Run2 data are also included, even though their poor performance could be due to insufficient sequencing depth.

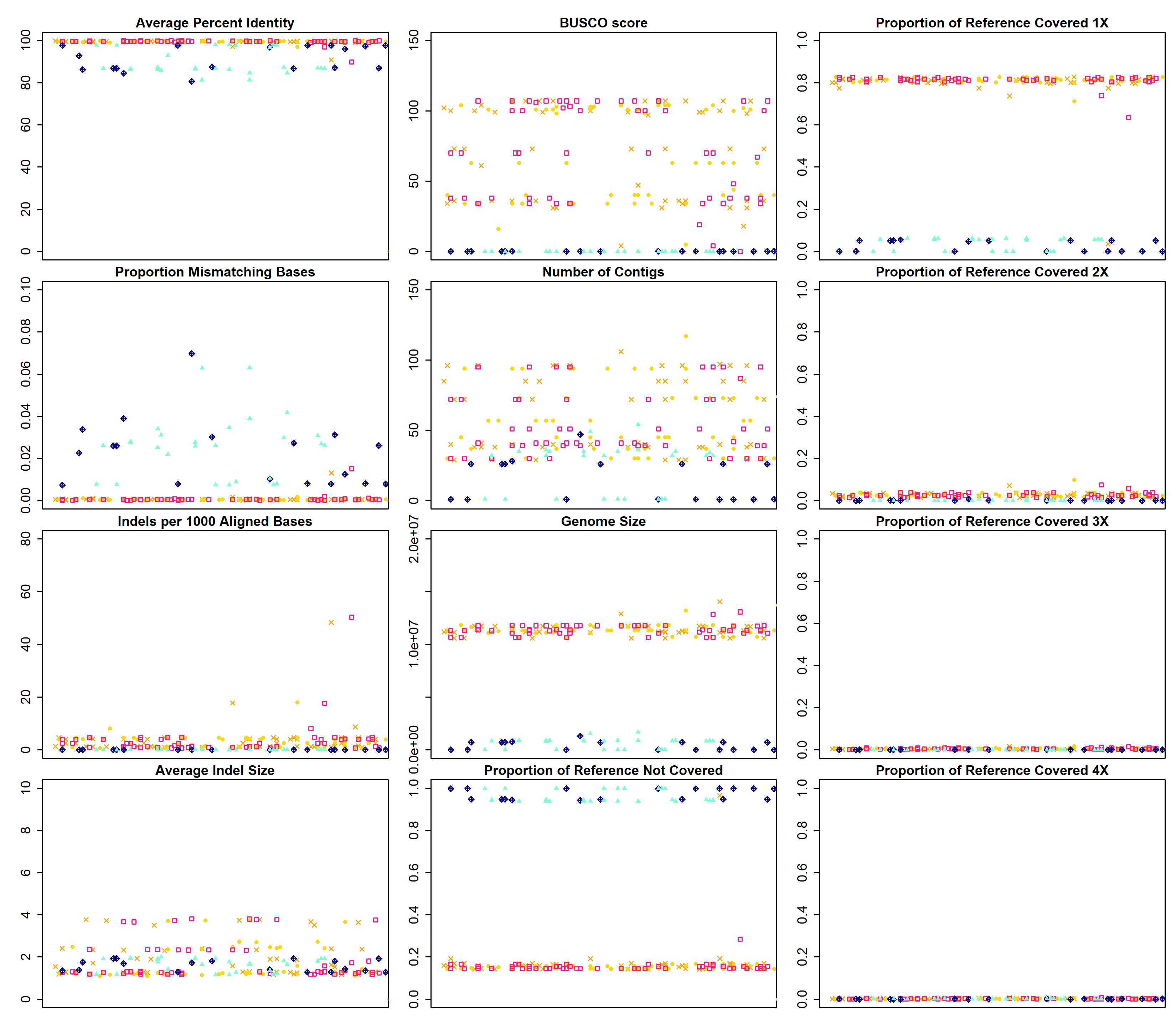

**Figure S4.** Performance metrics for pooled input and non-pooled input *Giardia* AWB assemblies. Orange X’s: non-pooled assemblies made from Run1_0157 reads, gold circles: pooled assemblies made from all reads from both Run1 and Run2 (0150, 0157, 2331, and 2338 reads), pink squares: pooled assemblies made from Run1 data only (0150 and 0157 reads), dark blue diamonds: pooled assemblies made from Run2 data only (2331 and 2338 reads), and aquamarine triangles: non-pooled assemblies made from Run2_2338 reads. The title above each scatterplot denotes the metric being plotted on the y-axis. The x-axis has no units because the x-values are randomly assigned to spread out the data points for visualization purposes. All metrics were calculated as described in the main text Methods. Assemblies from Run2 data are also included, even though their poor performance could be due to insufficient sequencing depth.

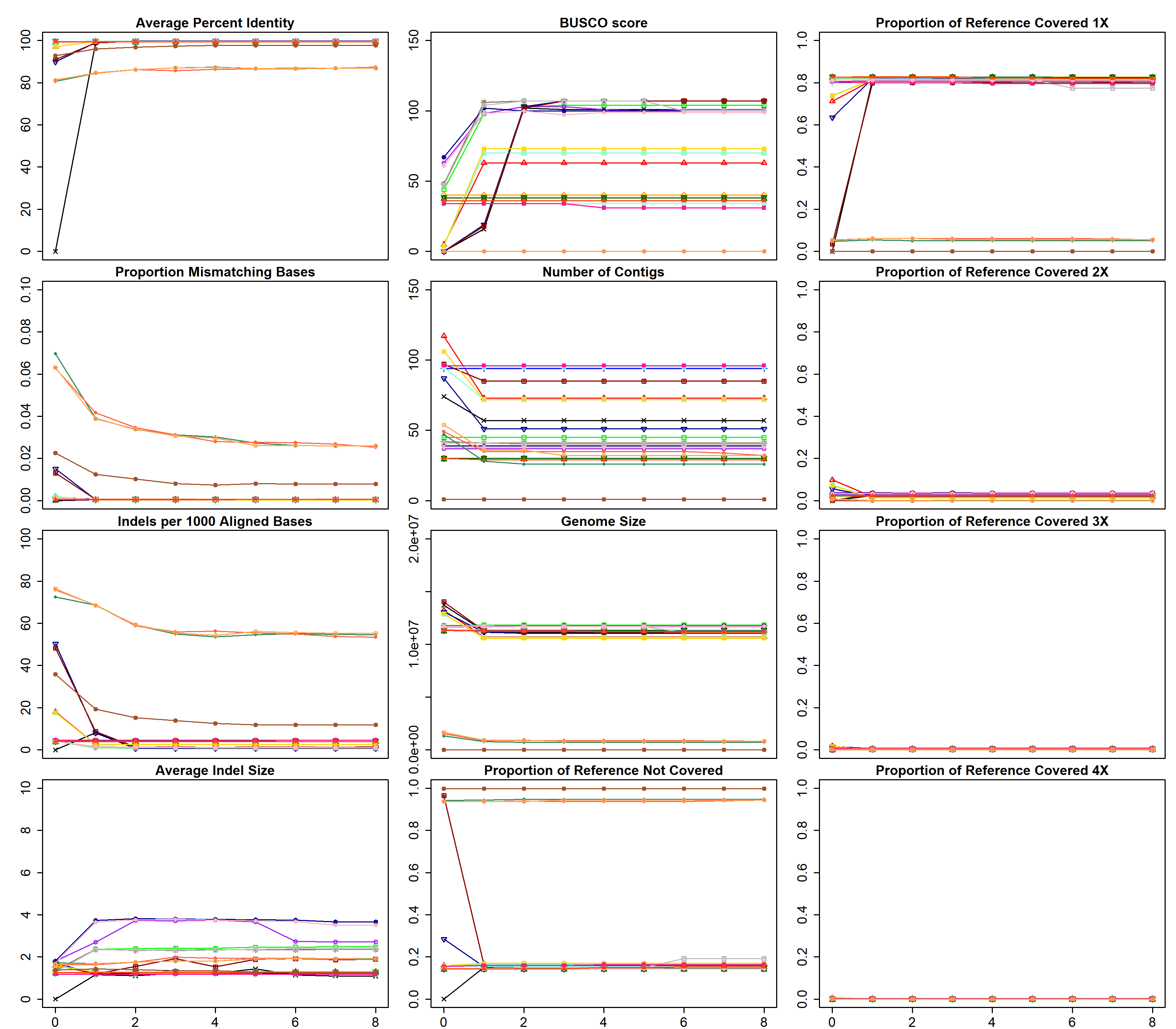

**Figure S5.** Performance metrics for polished sets of *Giardia* AWB assemblies. The title above each scatterplot denotes the metric being plotted on the y-axis. The x-axis denotes how many times the draft assembly has been polished. All metrics were calculated as described in the main text Methods. Assemblies from Run2 data are also included, even though their poor performance could be due to insufficient sequencing depth.

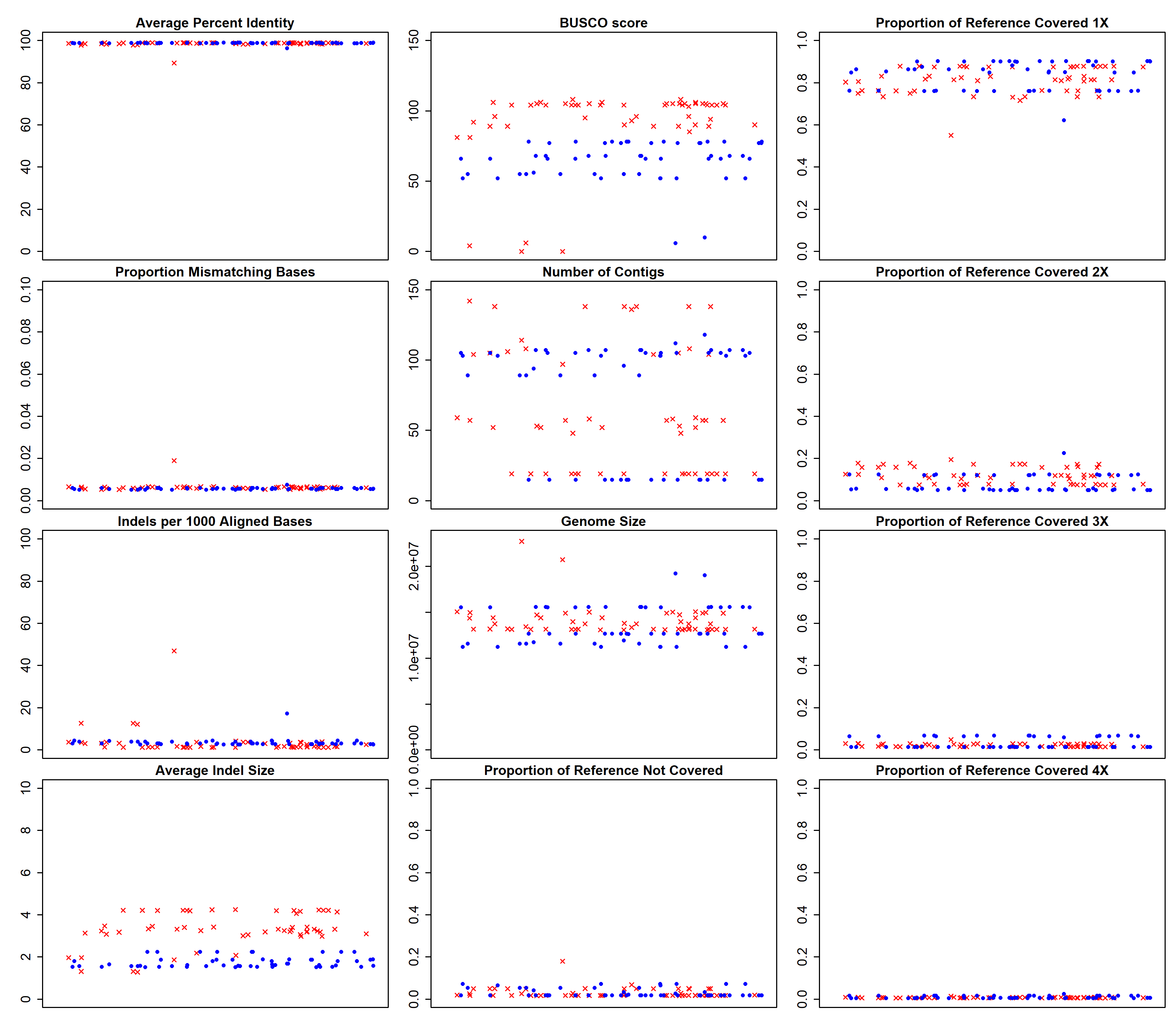

**Figure S6.** Performance metrics for all 1D and 1Dsq *Giardia* BGS assemblies. Red X’s denote all assemblies created from 1D reads and blue circles denote all assemblies created from 1Dsq reads. The title above each scatterplot denotes the metric being plotted on the y-axis. The x-axis has no units because the x-values are randomly assigned to spread out the data points for visualization purposes. All metrics were calculated as described in the main text Methods.

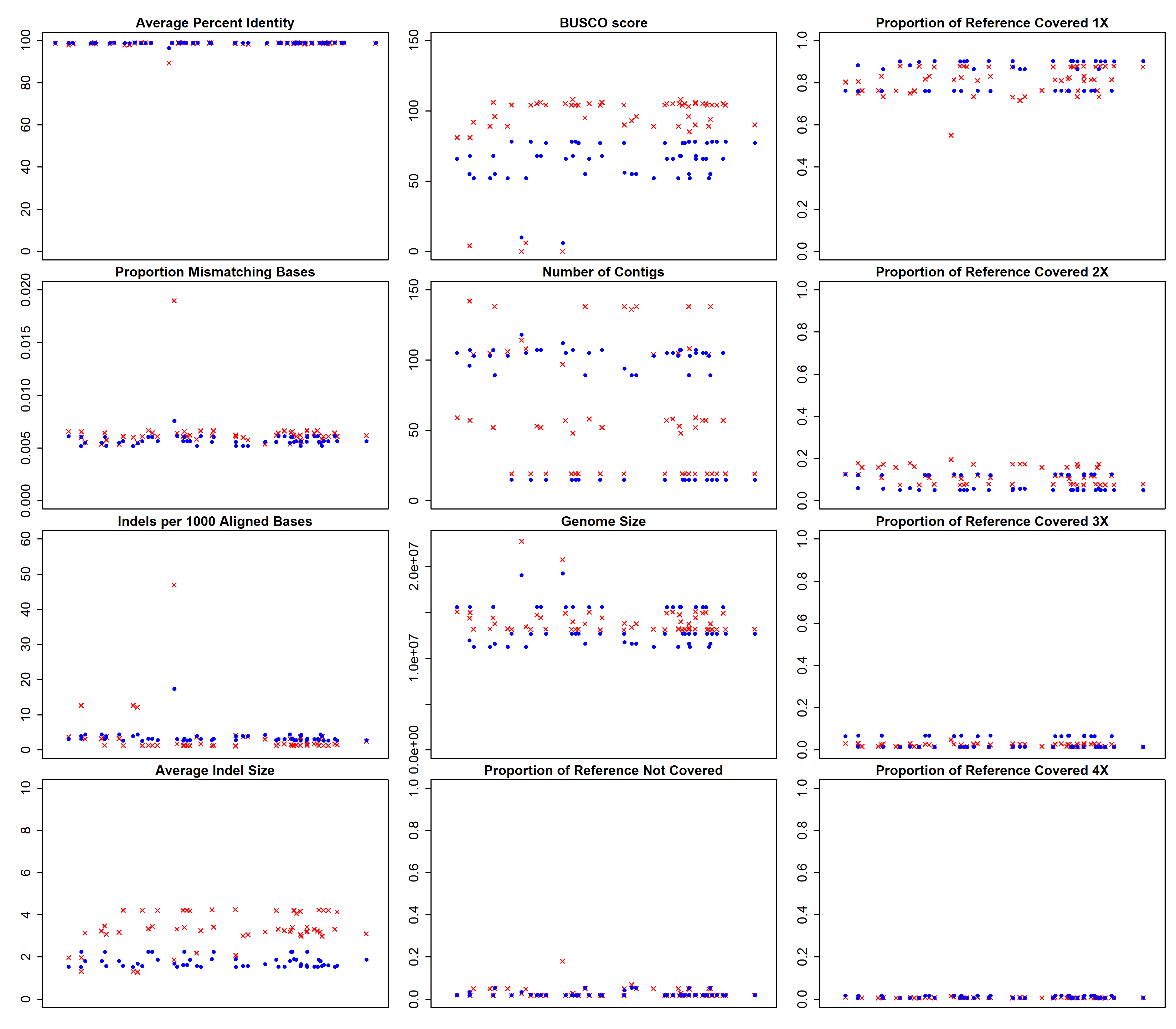

**Figure S7.** Performance metrics for corresponding 1D and 1Dsq *Giardia* BGS assembly pairs. All assemblies made from 1Dsq reads had a corresponding assembly made from 1D reads from the same inputs. Each pair was assigned the same random x-value so that corresponding 1D and 1Dsq assemblies would stack on top of each other in each plot. The additional assemblies from 1D reads are not shown. Red X’s denote all assemblies created from 1D reads and blue circles denote all assemblies created from 1Dsq reads. The title above each scatterplot denotes the metric being plotted on the y-axis. The x-axis has no units because the x-values are randomly assigned to spread out the data points for visualization purposes. All metrics were calculated as described in the main text Methods.

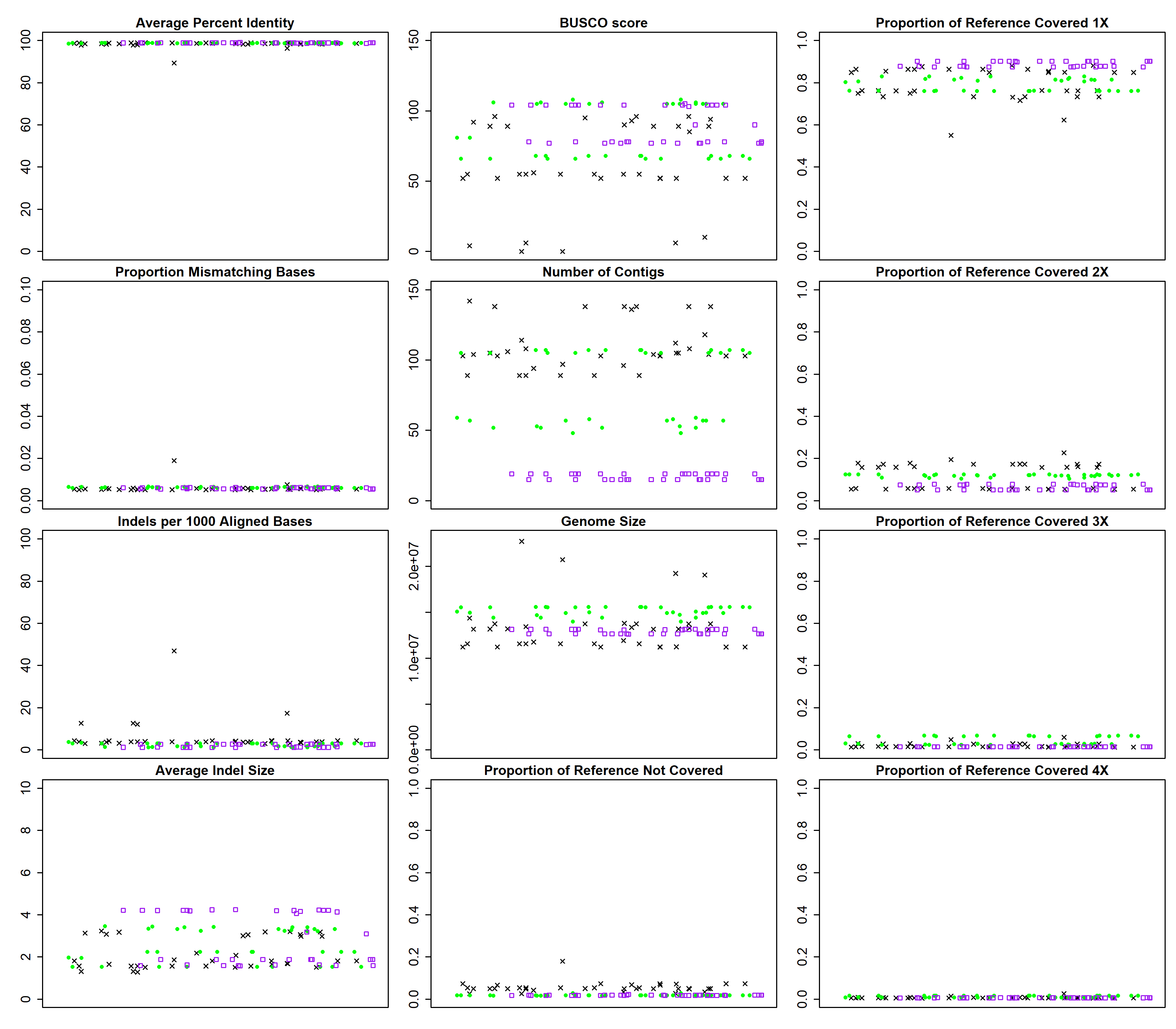

**Figure S8.** Performance metrics for all *Giardia* BGS assemblies, separated by assembly program. Black X’s denote all Abruijn assemblies, green circles denote all Canu assemblies, and purple squares denote all SMARTdenovo assemblies. The title above each scatterplot denotes the metric being plotted on the y-axis. The x-axis has no units because the x-values are randomly assigned to spread out the data points for visualization purposes. All metrics were calculated as described in the main text Methods.

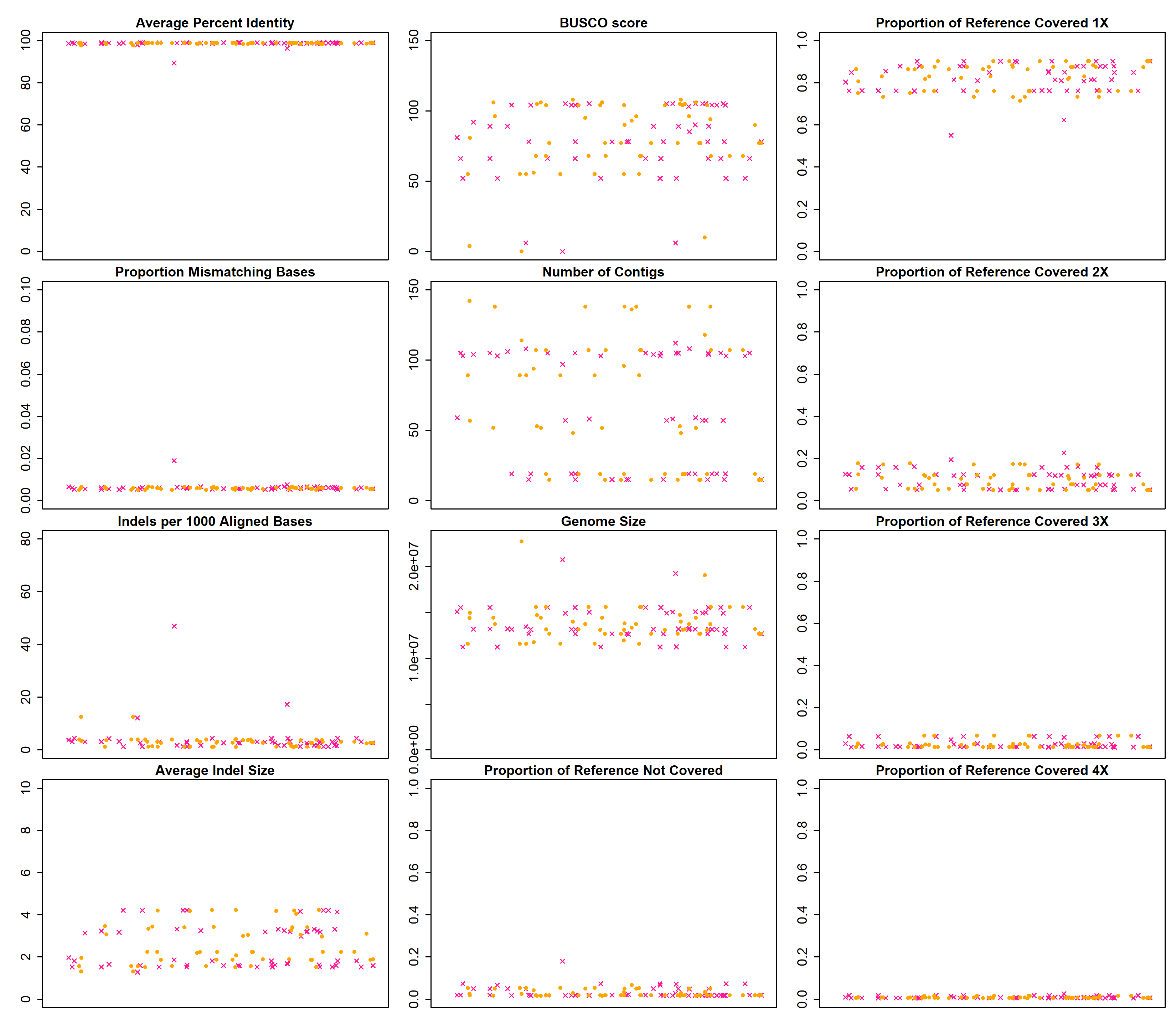

**Figure S9.** Performance metrics for pooled input and non-pooled input *Giardia* BGS assemblies. Pink X’s: pooled assemblies made from Run4 reads (2237, 2244), orange circles: non-pooled assemblies made from Run1_2244 reads. The title above each scatterplot denotes the metric being plotted on the y-axis. The x-axis has no units because the x-values are randomly assigned to spread out the data points for visualization purposes. All metrics were calculated as described in the main text Methods.

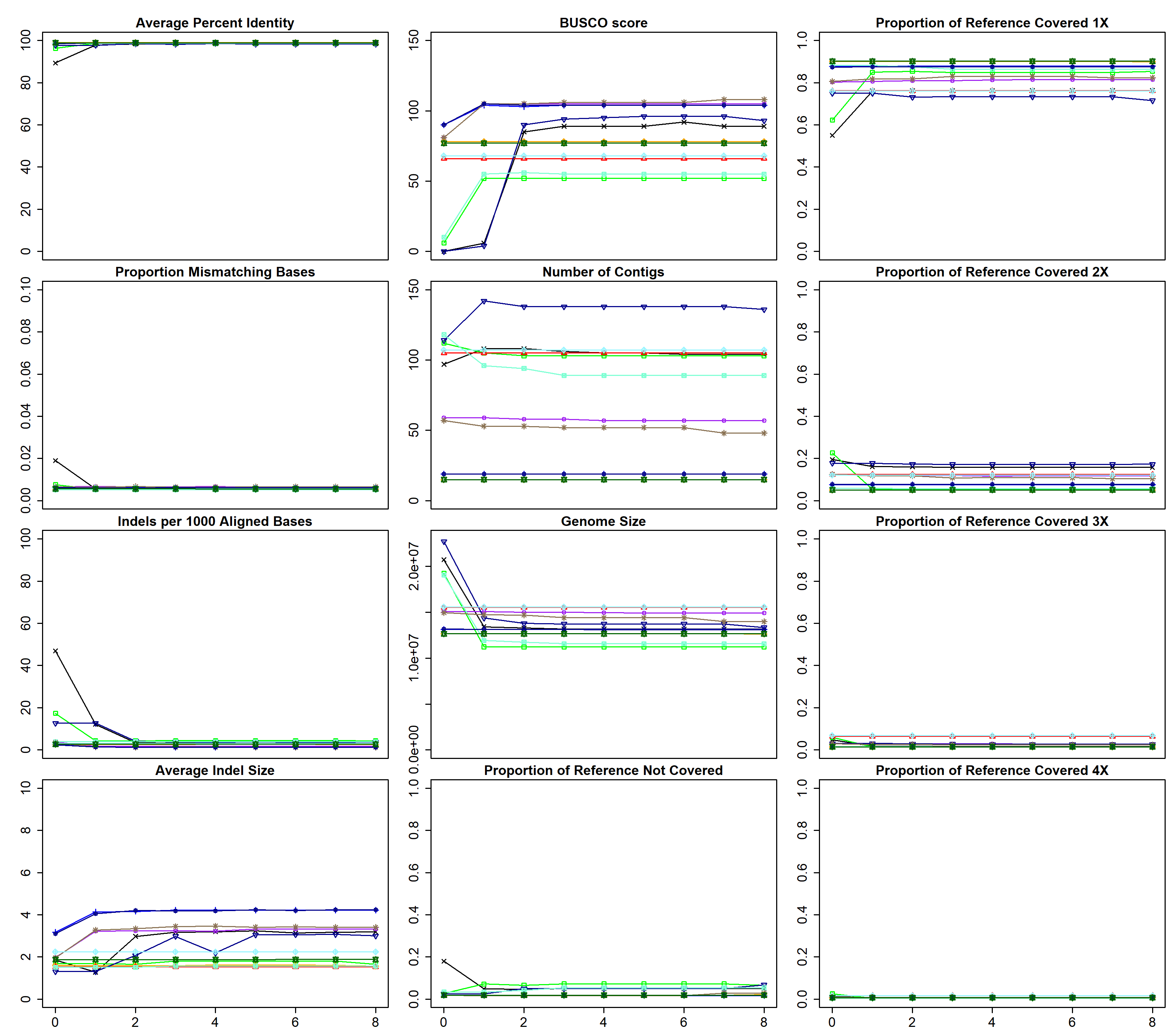

**Figure S10.** Performance metrics for polished sets of *Giardia* BGS assemblies. The title above each scatterplot denotes the metric being plotted on the y-axis. The x-axis denotes how many times the draft assembly has been polished. All metrics were calculated as described in the main text Methods.
