## Supplemental Text for "Nanopore sequencing of *Giardia* reveals widespread intra-isolate structural variation"

**Supplementary Discussion**

*Long read assemblies*

Upon initial inspection, the averages and standard deviations of the evaluation metrics for the assemblies generated from 1D vs 1Dsq input reads would suggest no difference between the two (Supplementary Table S3 – S4). The standard deviations also indicate performances of the 1D assemblies are more variable (Supplementary Table S3 – S4). However, plotting the values (Supplementary Figs. S1, S6), suggests that when the 1D assemblies perform well they often out-perform the 1Dsq assemblies, but they are also much more variable. It is also worth noting that most of the AWB 1D assemblies with poor performance in any of the evaluation metrics come from using only Run2 data (AWB_2331 and/or AWB_2338) and may be performing poorly due to insufficient sequencing depth (Supplementary Table 1). Since every assembly constructed from 1Dsq input reads has a corresponding assembly constructed from 1D input reads (Supplementary Table S15), to further examine the relationship between using 1D vs 1Dsq input reads, the 1D vs 1Dsq input pairs were plotted together (Supplementary Figs. S2, S7). The new plots show that while the 1D assemblies are often more variable than the 1Dsq assemblies, the 1D assemblies generally out-perform the 1Dsq assemblies, especially in the average percent identity, number of indels per 1000 aligned bases, BUSCO score, number of contigs, and genome size metrics (Supplementary Figs. S2, S7).

When examining the effects of pooling or not pooling runs for the same organism, the most obvious difference is between AWB assemblies generated from solely Run2 data (AWB_2331 and AWB_2338) and assemblies that include Run1 data (AWB_0150 and AWB_0157) (Supplementary Table S1, Supplementary Figs. S4, S9). Since this difference may be caused by the much smaller number of reads in Run2, informative comparisons of the effects of pooling or not pooling runs are AWB_0157 assemblies compared to AWB_0150_0157 and AWB_0150_0157_2331_2338 assemblies or AWB_2338 assemblies compared to AWB_2331_2338 assemblies. Among the assemblies that used Run1 data, no input combination produced values significantly different from the others for any metric examined (Supplementary Table S5) and examination of the plotted values shows no clear patterns for any metric (Supplementary Fig. S4), suggesting pooling or not pooling the input data had no effect. Similarly, the BGS assemblies and the AWB pooled and non-pooled Run2 assemblies did not have significantly different values for any metric (Supplementary Table S5, S6), nor did any clear pattern emerge when plotting the values (Supplementary Fig. S4, S9). Taken together these results suggest pooling or not pooling input data for the same organism has no significant effect on the final assembly once adequate genome coverage is achieved (though the exact cut-off for “adequate coverage” was not determined here). For runs with low read counts however, pooling runs can improve the final assembly, as was the case here for AWB_0150_0157_2331_2338 assemblies compared to AWB_2331_2338 assemblies.

Among the three assemblers tested, the SMARTdenovo assemblies showed the lowest variability in all metrics except average indel size (Main text Fig. 1). Moreover, the SMARTdenovo assemblies had the highest average values for average percent identity, BUSCO score, and proportion of reference covered 1X (where higher values indicate better performance) (Supplementary Tables S7, S8). They also had the lowest average value for proportion of reference not covered and an average genome size value that is closest to the size of the reference genome (Supplementary Tables S7, S8). Additionally, seven (2 AWB, 5 BGS) of the top performing assemblies in Supplementary Table S2 are SMARTdenovo assemblies. Plotting the SMARTdenovo assembly values for each metric also showed consistently strong performance in all metrics except average indel size (Supplementary Figs. S3, S8). The Abruijn assemblies show the greatest variability in all metrics except average indel size, number of contigs, and genome size (Supplementary Tables S7, S8). They had the lowest average indel size and the lowest variability in average indel size (Supplementary Tables S7, S8). Despite thirteen of the top performing assemblies in Supplementary Table S2 being Abruijn assemblies, plotting the Abruijn assembly values for each metric showed highly variable performance consistent with the averages and standard deviations in Supplementary Tables S7, S8 (Supplementary Figs. S3, S8). Finally, the Canu assemblies generally performed somewhere between the SMARTdenovo and Abruijn assemblies (Supplementary Tables S7, S8). Notably, all of the AWB assemblies with poor performance in the number of contigs metric were Canu assemblies (Supplementary Figs. S3, S8) and these were all generated from Run2 data only (Supplementary Table 1), suggesting that Canu is particularly sensitive to low coverage compared to the other assemblers.

The effects of genome polishing on each of the assembly evaluation metrics are shown in Supplementary Figs. S5, S10. For all metrics the biggest changes occur after the first or second round of polishing, after which they remain relatively consistent. For average percent identity, unpolished genomes that perform well show no significant change with polishing. For the unpolished genomes that do not perform well, polishing improves average percent identity up to about four or five rounds of polishing, after which average percent identity levels off (Supplementary Figs. S5, S10). The same trend can be seen for the proportion of mismatching bases metric where genomes that show an improvement with more than one round of polishing level off after around four or five rounds of polishing. Interestingly, the genomes that showed the biggest changes in the average indel size metric showed a decrease in performance with increasing polishing, though this decrease levelled off after around two or three rounds of polishing (Supplementary Figs. S5, S10). Finally, most improvements to the BUSCO score metric were seen after two rounds of polishing, after which values levelled off or occasionally decreased as polishing rounds reached six or higher (Supplementary Figs. S5, S10). Overall these results suggest four to five rounds of polishing will increase or not affect the performance of assemblies in all the metrics used here except average indel size.
